# A genetically encoded probe for imaging HA-tagged protein translation, localization, and dynamics in living cells and animals

**DOI:** 10.1101/474668

**Authors:** Ning Zhao, Kouta Kamijo, Philip D. Fox, Haruka Oda, Tatsuya Morisaki, Yuko Sato, Hiroshi Kimura, Timothy J. Stasevich

## Abstract

To expand the toolbox of imaging in living cells, we have engineered a new single chain variable fragment (scFv) that binds the classic linear HA epitope with high affinity and specificity *in vivo*. The resulting probe, which we call the HA frankenbody, is capable of lighting up in multiple colors HA-tagged nuclear, cytoplasmic, and membrane proteins in diverse living cell types. The HA frankenbody also enables state-of-the-art single-molecule experiments, which we demonstrate by tracking single mRNA translation dynamics in living U2OS cells and neurons. In combination with the SunTag, we track two mRNA species simultaneously to demonstrate comparative single-molecule studies of translation can now be done with genetically encoded tools alone. Finally, we use the HA frankenbody to precisely quantify the expression of HA tagged proteins in developing zebrafish embryos. The versatility of the HA frankenbody makes it a powerful new tool for imaging protein dynamics *in vivo*.

**One-sentence summary:** A genetically encodable intracellular single-chain variable fragment that selectively binds the HA epitope (YPYDVPDYA) with high affinity in living cells and organisms can be used to quantify HA-tagged protein translation, localization, and dynamics.

## INTRODUCTION

Live-cell imaging is critical for tracking the dynamics of cell signaling. The discovery and development of the green fluorescent protein (GFP), for example, has literally revolutionized the field of cell biology^1,2^. GFP can be genetically fused to a protein of interest (POI) to light it up selectively and track its expression and localization in living cells and organisms. While incredibly powerful, GFP-tagging has several limitations that have made it difficult to image the full lifecycles of proteins. First, long fluorophore maturation times prevent co-translational imaging of GFP-tagged nascent peptide chains^3,4^. By the time the GFP tag folds, matures and lights up, translation of the nascent peptide chain is long over. Similarly, slow GFP maturation times have made it difficult to image short-lived transcription factors that are critical for development and embryogenesis^5^. Again, before the GFP tag has time to light up, the transcription factor may already be degraded. Second, GFP fusion tags cannot discriminate post-translational modifications (PTM) to proteins^6^, such as acetylation, methylation and phosphorylation, nor can they discriminate protein conformational changes^7^. Without the ability to directly image these important protein subpopulations, their unique functionality is difficult to quantify and assess. Third, GFP fusion tags are relatively large, permanently attached, and dim. It is therefore difficult to detect and/or amplify fluorescence signal. This severely limits the length of time a single tagged protein can be tracked in a living cell before the protein is either photobleached or the cell is photodamaged.

To address these limitations of GFP, an alternative live-cell imaging methodology has recently emerged that uses antibody-based probes^8^. In this methodology, probes built from antibodies, such as antigen binding fragments (Fabs)^9^, single-chain variable fragments (scFvs)^10–12^ and camelid nanobodies^13–16^, are conjugated or genetically fused with mature fluorophores. When expressed or loaded into cells, the probes dynamically bind and light up epitopes within POIs as soon as the epitopes are accessible. With this emerging methodology, it is possible to visualize and quantify the co-translational dynamics of nascent peptide chains^17–21^, capture the dynamics and localization of short-lived transcription factor dynamics in living embryos^5^, track single molecules for extended periods of time^22,23^, and selectively track the spatiotemporal dynamics of PTMs^24^ and specific protein conformational changes^7^.

While there is great potential for antibody-based probes in live-cell imaging, so far only a small handful have been developed and tested. Arguably the most straight-forward probes to develop and test are Fab, since they can be digested from commercially available antibodies and conjugated with dyes using standard kits. For example, we have generated a variety of complementary Fab to track both endogenous histone modifications and single mRNA translation dynamics^6,17,24,25^. Unfortunately, Fab have not been widely adopted, in large part because of the difficulties associated with loading them into living systems. While robust adherent cells, such as Hela or U2OS, can be easily bead loaded in mass^26^, more sensitive cell types, including neurons and embryonic stem cells, have proven refractive to most loading procedures. Additionally, Fab are expensive to work with, typically requiring milligrams of purified full-length antibody as starting material. Fab may also change considerably from batch to batch, which can lead to unwanted variability between experiments that complicates downstream analyses.

Given the drawbacks of Fab, genetically encoded probes are an attractive alternative. Since these probes can be integrated into plasmids, they can easily be distributed and cell lines and/or transgenic organisms can be generated that stably express the probes, all without the worry of batch-to-batch variability. The only downside of genetically encoded antibody-based probes is they are not straightforward to develop. Both scFv and camelid nanobodies require a large initial investment, as either existing hybridomas or immunized animals are necessary to get the sequences of individual antibody chains. Worse, even after specific sequences are determined, there is a good chance that antibody-based probe derived from the sequences will not fold and function properly *in vivo*. The problem is that antibodies have evolved to be secreted from cells, so their folding and maturation is more often than not disrupted when expressed within the reduced intracellular environment. This leads to low-expression and aggregation inside living cells. For example, in our experience developing scFv against endogenous histone modifications (i.e. mintbodies), only around 5% of antibody heavy and light-chain sequences properly fold when integrated into an scFv framework and expressed within living mammalian cells. Camelid nanobodies also suffer from misfolding, instability and aggregation when expressed inside living cells. Thus, when it comes to antibody-based probes for imaging, sequence information is usually not enough. Instead, extensive protein engineering, directed evolution, and mutagenesis are typically needed to generate an ideal probe that functions well both inside and outside living cells.

A case in point is the SunTag scFv, the only genetically encoded antibody-based probe capable of binding a small epitope co-translationally in living cells. The SunTag scFv binds a 19 aa epitope (EELLSKNYHLENEVARLKK) that is repeated 24 times within a single SunTag. As multiple scFvs tightly bind the SunTag co-translationally, fluorescence signal from individual tagged POIs can be greatly amplified, enabling both single mRNA translation imaging and long-term single molecule tracking *in vivo*^18–21,27,28^. The SunTag imaging technology was developed over many years, starting with work in 1998 by the Plϋckthun lab. Briefly, after generating antibodies from immunized mice, ribosome display was used to affinity select high-performance scFv variants^29^. With the aid of crystal structures and computational design, the complementarity determining regions (CDRs or loops) of these scFv were grafted onto an in-vitro optimized hyper-stable scFv scaffold mutagenized at about 20 individual residues to minimize grafting mismatch^30^. This led to a stable version 1 probe that was later tested in 2014 by Tanenbaum et al to stain mitochondria in living cells^23^. The probe was further optimized via the addition of stabilizing sfGFP and GB1 domain to eliminate aggregation at higher expression levels and the original epitope was optimized to version 4 via directed mutagenesis.

The large amount of work required to develop the SunTag imaging technology highlights the difficulty of generating new antibody-based probes that are suitable for demanding live-cell imaging applications. To confront this problem, we here develop an alternative strategy for developing new scFvs for live-cell imaging. To bypass many of the difficulties associated with probe development, we begin with a diverse set of scFv scaffolds that have already been proven to fold properly and function within the reduced cytoplasm of living cells. Onto these scaffolds, we loop graft all six CDRs from an epitope-specific antibody^31^. Depending on the compatibility of the scaffold and CDRs, this produces a hybrid scFv that retains the folding stability of the scaffold, while acquiring the binding specificity of the grafted CDRs.

To demonstrate the efficiency of our approach, we use it to generate two new hybrid scFvs (with a 40% success rate) that bind to the classic linear HA epitope (YPYDVPDYA)^32^ in living cells and organisms. We extensively tested one of these hybrid scFvs, which we call the “HA frankenbody,” by using it to label in multiple colors a variety of proteins in diverse live-cell environments. In all cases we tested, the HA frankenbody binds its target HA-epitope with high affinity and selectivity, making it suitable for a wide range of imaging applications. This makes the HA frankenbody a powerful new imaging tool to study complex protein dynamics in living systems with high spatiotemporal resolution. We anticipate that as additional antibody sequences become available, our general strategy to develop frankenbodies will become easier and more efficient, leading to a complementary set of scFvs for highly multiplexed imaging of the full lifecycle of proteins *in vitro* and *in vivo*.

## RESULTS

### Design strategy and initial screening of frankenbodies

We engineered the HA-frankenbody from six complementarity determining regions (CDRs, or loops) within the heavy and light chains of a published anti-HA scFv (parental full-length antibody: 12CA5)^33,34^ (Fig. 1A). On its own, this wildtype anti-HA scFv (wtHA-scFv) does not fold properly in the reduced intracellular environment, and therefore displays little to no affinity for HA epitopes in living human U2OS cells^33^. We figured we could address this folding issue by grafting the CDRs onto more stable and sequence similar scFv scaffolds (Fig. 1A). To test this, we selected five scFv scaffolds that have already been successfully used for live-cell imaging purposes and that have a wide range of sequence identity compared to wtHA-scFv. In particular, the sequences of the heavy chain variable regions (VH) were 47-89% identical, while the sequences of the light chain variable regions (VL) were 50-67% identical. The five scaffolds we chose included (1) an scFv that specifically binds histone H4 mono-methylated at Lysine 20 (H4K20me; 15F11)^35^; (2) an H3K9ac specific scFv (13C7)^36^; (3) an H420me2-specific scFv (2E2, unpublished); (4) a SunTag-specific scFv^23^; and (5) a bone Gla protein (BGP) specific scFv (KTM219)^33^. Among these scaffolds, 15F11 and 2E2 have the greatest sequence identity compared to the wtscFv-HA (Fig. 1B). We therefore hypothesized there would be a higher chance of grafting success with either of these scaffolds.

**Figure 1.**
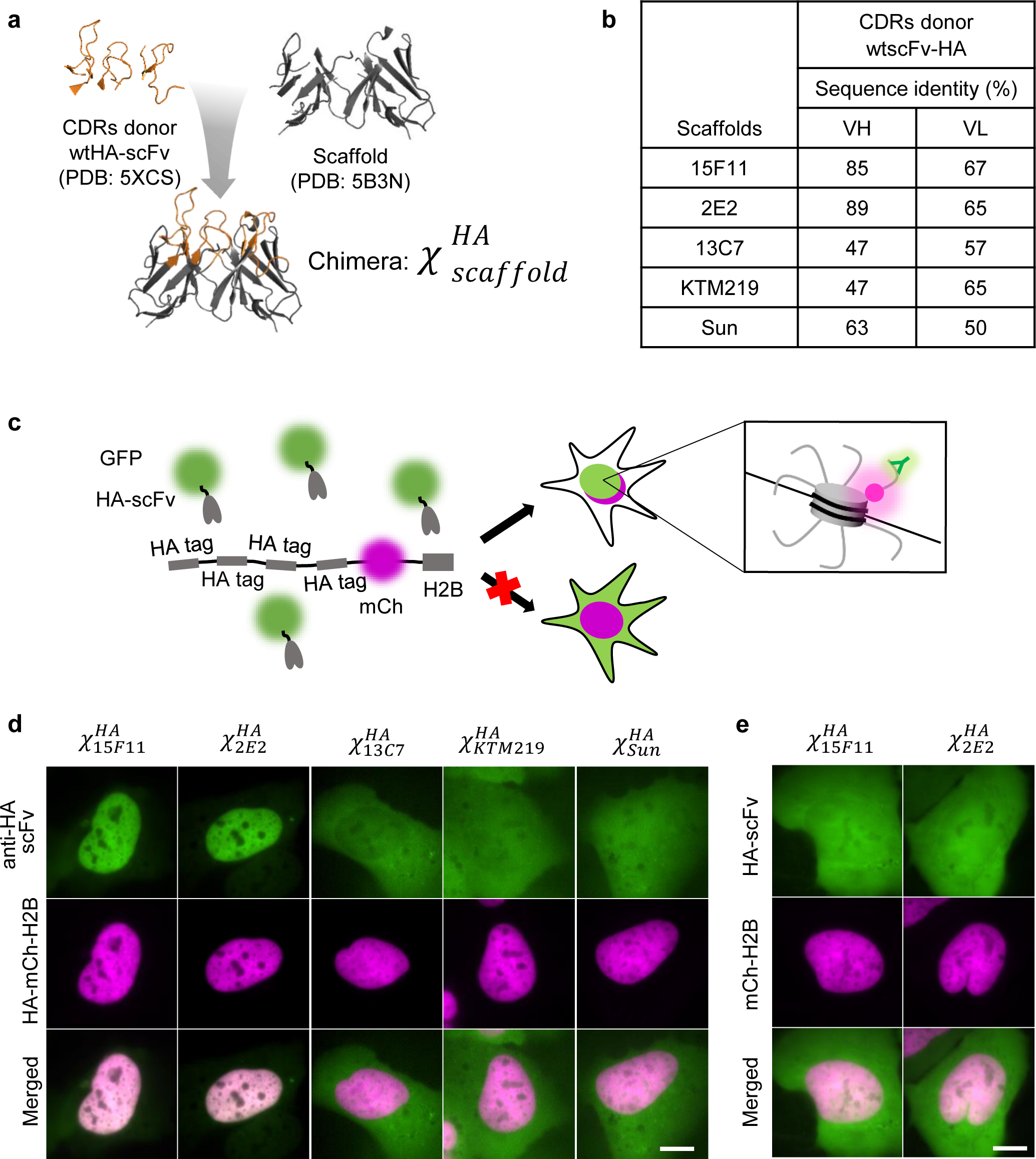
Design strategy and initial screening of frankenbodies. (a) A cartoon schematic showing how to design a chimeric anti-HA scFv using wtHA-scFv CDRs and stable scFv scaffolds. (b) Sequence identity analysis of the wtHA-scFv with five intracellular scFv scaffolds. (c) A cartoon showing how to screen the five chimeric anti-HA scFvs in living U2OS cells. (d) Initial screening results showing the respective localization of the five chimeric anti-HA scFvs in living U2OS cells co-expressing HA-tagged histone H2B (chimeric anti-HA scFv, green; HA-mCh-H2B, magenta). (e) Control results showing the respective localization of 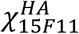 and 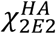 in living cells lacking HA-tagged histone H2B (chimeric anti-HA scFv, green; mCh-H2B, magenta). Scale bars: 10µm.

To verify our hypothesis, we grafted the anti-HA scFv CDR loops onto our five chosen scFv scaffolds. We refer to the resulting five chimeric scFvs as 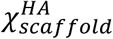. For example, 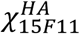 specifies the chimeric scFv that was generated by loop grafting the anti-HA CDRs onto the 15F11 scFv scaffold. To screen our chimeras, we fused each with the green fluorescent protein mEGFP and co-transfected each of the resulting plasmids into U2OS cells, together with a plasmid encoding 4×HA-tagged histone H2B fused to the red fluorescent protein mCherry (HA-mCh-H2B). If a chimeric scFv binds to the HA epitope in living cells, it should co-localize with the HA-tagged H2B in the nucleus, as shown in Fig. 1C. Live-cell imaging revealed 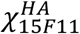 and 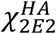 were superior, displaying little to no misfolding and/or aggregation, strong expression, and excellent co-localization with H2B in the nucleus. In contrast, the other three scFvs did not show any co-localization signal (Fig. 1D). Moreover, in control cells lacking HA tags, both 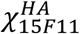 and 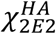 displayed uniform expression (Fig. 1E), indicative of free diffusion without non-specific binding. According to our screen, both 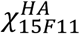 and 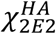 worked equivalently well in living cells. We chose the 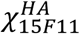 variant for additional screening, which we herein refer to as the “HA frankenbody” due to its construction via grafting.

### Multicolor labeling of HA-tagged proteins in diverse intracellular environments

We tested the HA frankenbody in a variety of different settings. First, as we already demonstrated in the initial screen, the HA frankenbody colocalizes with HA-tagged histone H2B in the nucleus of living U2OS cells (Fig. 2A). This demonstrates frankenbodies can pass through the nuclear pore and bind target nuclear proteins. We next wanted to test if the HA frankenbody can work equally well in the cell cytoplasm, another reducing environment that can interfere with intradomain disulfide bond formation^30^. We tested this by creating a new target plasmid encoding the cytoplasmic protein β-actin fused with a 4×HA-tag and mCherry (HA-mCh-β-actin). When this plasmid was expressed in cells, co-expressed frankenbodies again took on the distinct localization pattern of their targets, in this case colocalizing with HA-mCh-β-actin along filamentous actin fibers (Fig. 2B). We therefore conclude that both nuclear and cytoplasmic HA-tagged proteins can be selectively labeled with high efficiency by the HA frankenbody in living human cells.

**Figure 2.**
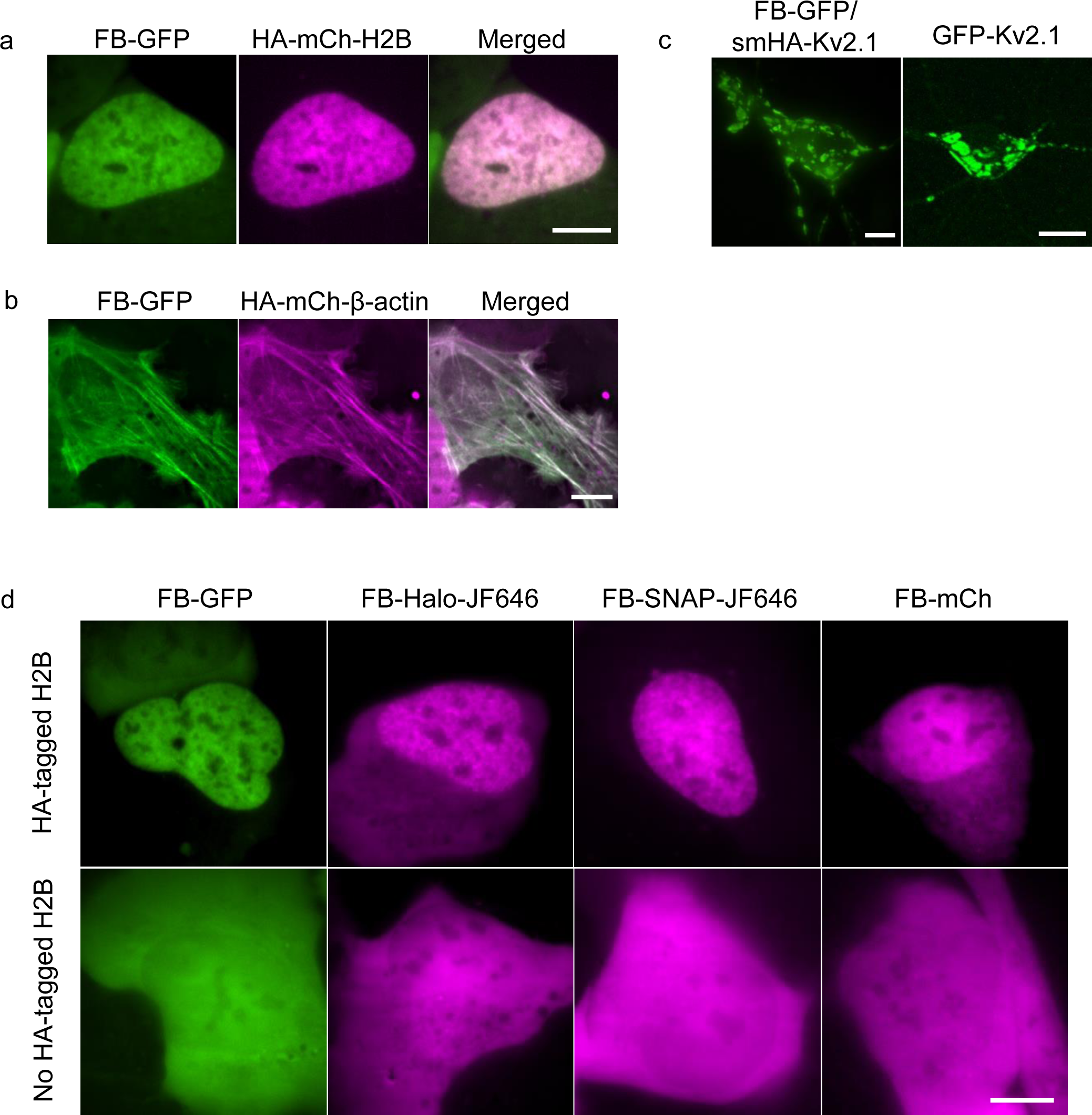
Multicolor labeling of HA-tagged proteins in diverse intracellular environments. (a) Frankenbody (FB-GFP; green) labels an HA-tagged nuclear protein, histone H2B (HA-mCh-H2B; magenta), in living U2OS cells. (b) FB-GFP (green) labels an HA-tagged cytoplasmic protein, β-actin (HA-mCh-β-actin; magenta), in living U2OS cells. (c) FB-GFP labels an HA-tagged membrane protein, Kv2.1 (left), showing similar localization as GFP-Kv2.1 (right) in living neurons. (d) FB fused to multiple fluorescent fusion proteins (top row: GFP, HaloTag-JF646, SNAP-tag-JF646 and mCherry) specifically labels HA-tagged nuclear protein H2B (HA-mCh-H2B). In all cases, cells lacking the HA-tag display relatively even FB fluorescence (bottom row). Scale bars, 10 µm.

To test if frankenbodies could also work in more sensitive cell types, we co-transfected living primary rat cortical neurons with the HA frankenbody (FB-GFP) and an HA-tagged transmembrane protein Kv2.1 (smHA-Kv2.1). In cortical and hippocampal neurons, GFP-Kv2.1 is localized to the plasma membrane, where it forms large (up to one micron in diameter) cell-surface clusters, providing a distinct localization pattern^37^ (Fig. 2C right panel). In cells expressing FB-GFP and smHA-Kv2.1, the HA frankenbody again took on the distinct localization pattern of its target (Fig. 2C left panel). In addition, the distinctive pattern could be seen for over a week after transient transfection. This demonstrates the HA frankenbody can selectively bind membrane proteins as well as cytoplasmic and nuclear proteins, depending on the presence of the HA tag. This also demonstrates continual expression of frankenbody does not detrimentally impact sensitive cells and, moreover, frankenbodies have exceptionally long half-lives when bound to their targets.

To ensure the HA frankenbody is as broadly applicable as possible, we wanted to test if it could tolerate different fluorescent protein fusion partners that might be needed in multicolor imaging applications. GFP and its derivatives are generally superior fusion partners because their high stability actually helps stabilize and solubilize the tagged protein. This was observed, for example, during the development of the SunTag scFv^23^. To test how well the HA frankenbody tolerates different tags, we fused it to mCherry, HaloTag^38^ and SNAP-tag^39^. Encouragingly, all three frankenbody constructs colocalized with HA-tagged H2B in the nucleus of living U2OS cells, similar to the original GFP-tagged frankenbody (Fig. 2D, upper). Furthermore, all three constructs displayed relatively diffuse localization patterns in cells lacking the HA-tag (Fig. 2D, lower). These data indicate the HA frankenbody can indeed tolerate different fusion partners, including green (GFP), red (mCherry), and far-red (SNAP-tag/HaloTag with far-red ligands) fluorescent proteins. Thus, the HA frankenbody can label HA-tagged proteins in a rainbow of colors in living cells.

### Immunostaining and Western blotting with purified recombinant frankenbody

We wondered if the HA frankenbody has the potential to replace costly anti-HA antibodies in traditional assays such as immunostaining and Western blots. To test this, we cloned the frankenbody gene fused with mEGFP and a hexahistidine tag into an *E. coli* expression vector, pET23b. We expressed the recombinant frankenbody and purified the soluble portion from *E. coli*. Using this purified fraction, we immunostained fixed cells expressing HA-tagged H2B and HA-tagged β-actin. Similar to our observations in living cells, the purified HA frankenbody beautifully stained both the HA-tagged nuclear and cytoplasmic proteins, but now with almost no observable background signal (Fig. 3A, B).

**Figure 3.**
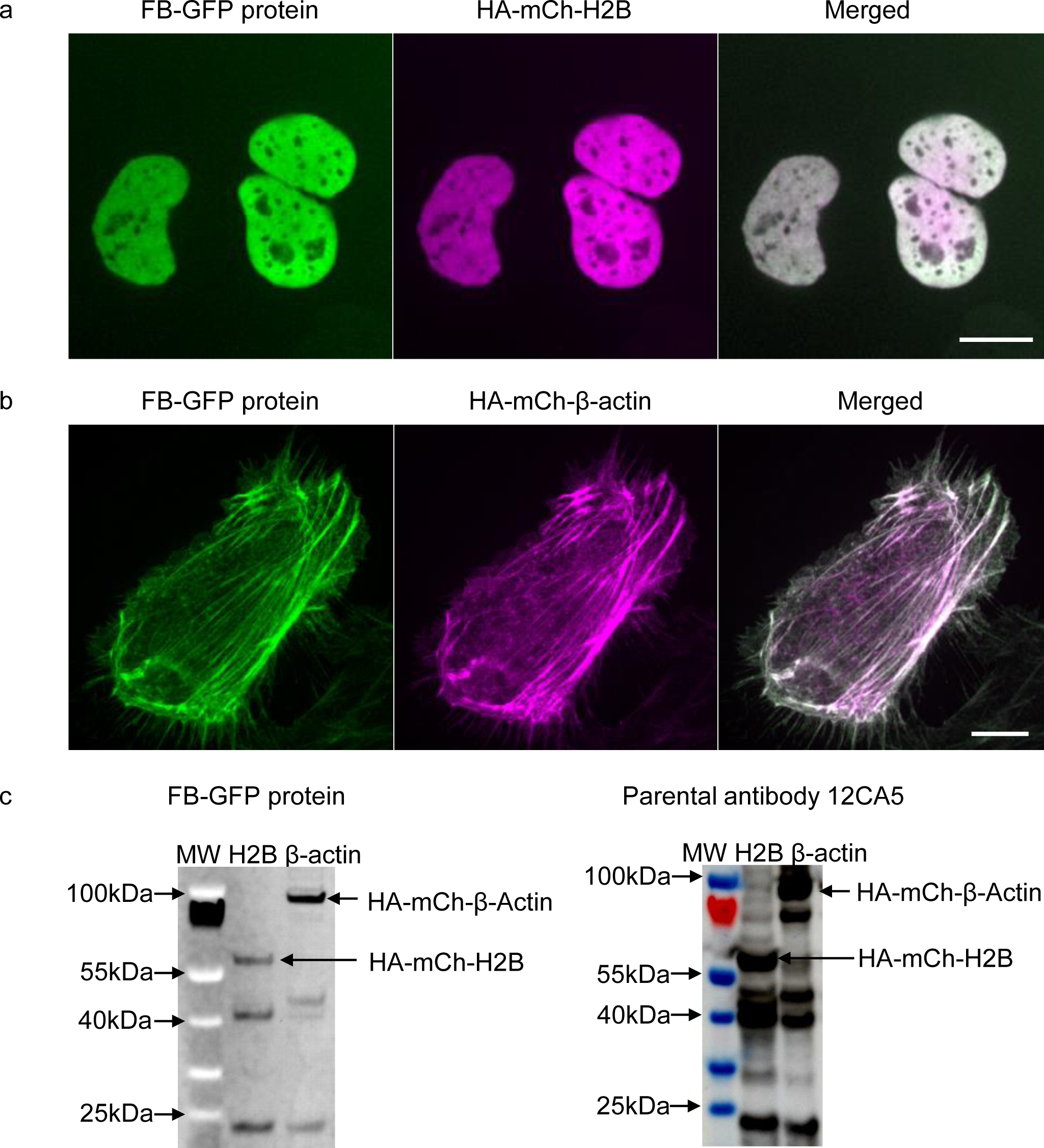
Immunostaining and Western blotting with purified recombinant frankenbody. Immunostaining in fixed U2OS cells with purified frankenbody (FB-GFP; green) of an HA-tagged (a) nuclear protein, histone H2B (HA-mCh-H2B; magenta) and (b) cytoplasmic protein, β-actin (HA-mCh-β-actin; magenta). (c) Western blot of HA-tagged H2B and β-actin. Left: purified FB-GFP (no secondary antibody) detected directly using GFP fluorescence; Right: parental anti-HA antibody 12CA5 detected with secondary anti-mouse antibody/HRP and the chemiluminescent substrate ECL. Scale bars, 10 µm.

We next tested the suitability of the HA frankenbody for Western blotting. For this, we harvested U2OS cells 24h after transiently transfecting either HA-tagged H2B or HA-tagged β-actin. In contrast to the parental 12CA5 full-length anti-HA antibody, which was stained using a secondary antibody and visualized via a sensitive chemiluminescent substrate, the frankenbody Western blot used the GFP signal alone for detection. Nevertheless, similar dark and sharp bands were seen on the frankenbody membrane as the 12CA5 membrane (Fig. 3C). Although several of the bands were dimmer than those seen using 12C15, we attributed the difference to signal amplification from the secondary antibody and the chemiluminescent substrate. In principle, a similar signal-to-noise could be attained using the GFP-tagged HA frankenbody with secondary antibodies against GFP. Together, our Western blot and immunostaining results strongly suggest the HA frankenbody can serve as a cost-effective replacement for full-length HA antibody in widely used *in vitro* applications.

### HA frankenbody specifically binds the HA epitope for minutes at a time in live cells

An ideal imaging probe binds its target with high affinity to maximize the fraction of target epitopes bound and thereby increase signal-to-noise. In general, a high bound fraction is established by a large ratio of probe:target binding off to on times; in other words, the time a probe remains bound to a target is ideally much longer than the time it takes a probe to bind a target. Although the latter depends sensitively on the concentrations of both target and probe, the former is a fixed biophysical parameter that is useful for planning and interpreting experiments. With this in mind, we set out to measure the length of time the HA frankenbody remains bound to the HA epitope in living cells.

To accurately measure the binding kinetics of HA frankenbody, we performed Fluorescence Recovery After Photobleaching (FRAP) experiments in cells co-expressing GFP-tagged HA frankenbody (FB-GFP) and HA-mCh-H2B. As H2B is bound to chromatin for hours at a time^40^, any recovery on the minutes timescale can be attributed to the turnover of frankenbody alone. Consistent with this, FRAP in the red HA-mCh-H2B channel displayed little or no recovery on the timescale of our experiment. In contrast, FRAP in the green FB-GFP channel slowly recovered (Fig. 4A). Quantitative analyses of FRAP curves revealed the half recovery time to be around 2~3 minutes (Fig. 4B). As a control, we repeated FRAP experiments in cells transfected with just the HA frankenbody (i.e. lacking HA epitopes). Here, the FRAP recovery lasted just seconds, consistent with little to no non-specific binding of the HA frankenbody (Fig. 4C). We therefore conclude that the majority of frankenbody bind target HA epitopes for minutes at a time in living cells. This is consistent with the high affinity binding (K_D_=13.4nM) we measured *in vitro* using surface plasma resonance (Fig. S2).

**Figure 4.**
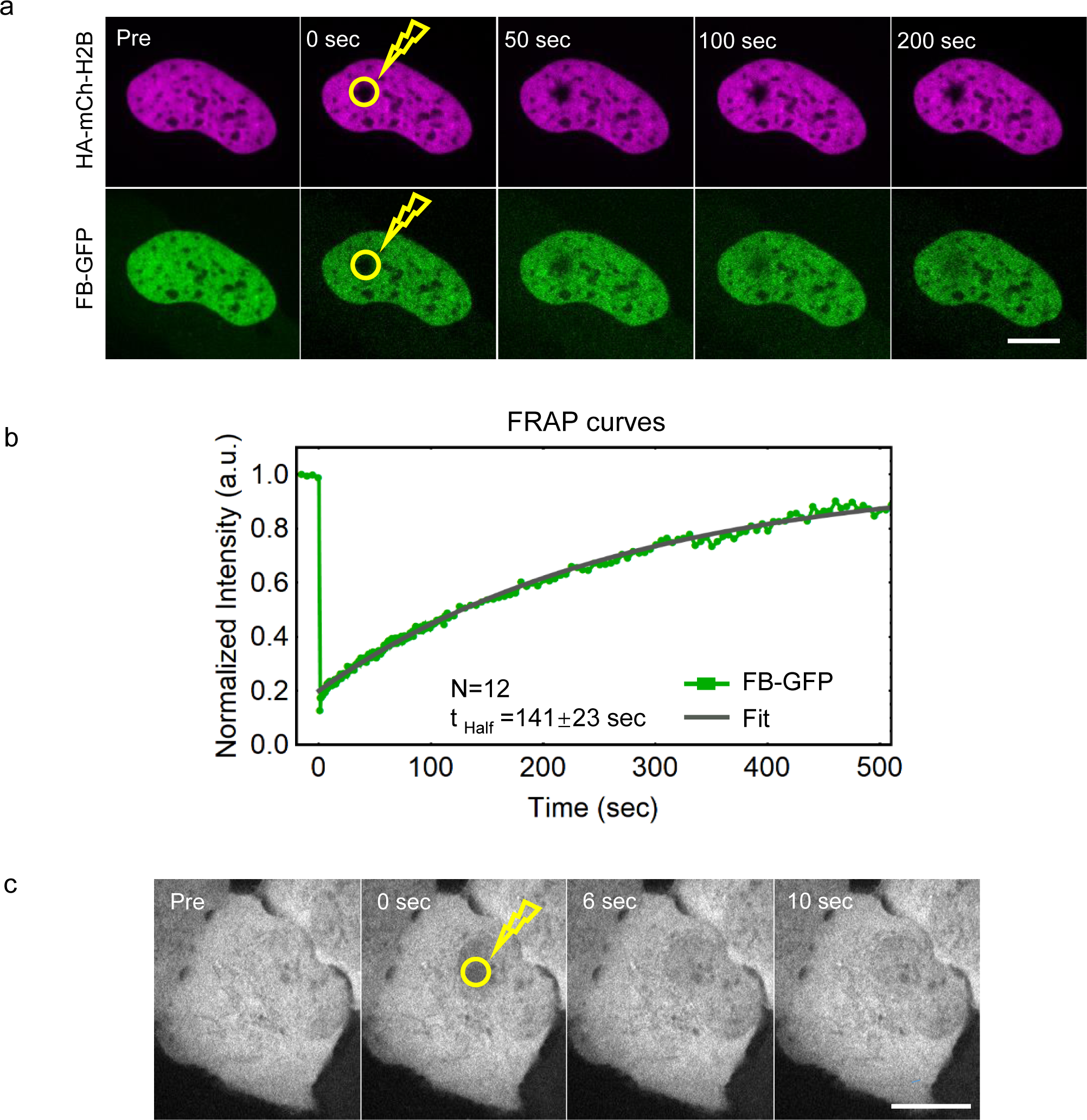
A frankenbody specifically binds the HA epitope for minutes at a time in live cells. (a) A representative FRAP experiment (yellow circle indicates bleach spot) showing fluorescence recovery in cells expressing frankenbody (FB-GFP; green) and target HA-mCh-H2B (magenta). (b) Quantification of FRAP data in a representative cell, along with a fitted curve. Fits from 12 cells reveal the half recovery time, *t*_*half*_, is 141±23 sec. (c) A representative FRAP experiment (yellow circle indicates bleach spot) in cells expressing FB-GFP only (i.e. cells lacking HA-tags) is complete in less than 10 seconds. Scale bars, 10 µm.

### Tracking single mRNA translation in living U2OS cells with the HA frankenbody

A major advantage of the HA frankenbody over other intrabodies is the small size and linearity of its epitope, just 9 aa in length. This means the epitope is quickly translated by the ribosome and becomes available for binding almost immediately. The HA frankenbody therefore has the potential to bind HA-tagged nascent peptides co-translationally, much like purified anti-HA antibody fragments are capable of^17^. By simply repeating the HA epitope multiple times within a tag, fluorescence can furthermore be amplified for sensitive single molecule tracking^22^.

To test the potential of HA frankenbody for imaging translation dynamics, we co-transfected a GFP-tagged version (FB-GFP) into U2OS cells together with our standard translation reporter. The reporter encodes a 10x HA spaghetti monster tag N-terminally fused to the nuclear protein KDM5B. In addition, the reporter contains a 24x MS2 stem loop repeat in the 3’ untranslated region to label and track single mRNA^17^ (Fig. 5A). A few hours after transfection, single mRNA (labeled by HaloTag-MS2 coat protein, MCP-HaloTag, and the JF646 HaloTag ligand^41^) could be seen diffusing throughout the cell cytoplasm. HA frankenbody comoved with many of these mRNA (Fig. 5B, Movie S1), indicative of active translation and similar to what is seen with anti-HA antibody fragments^17^. To prove these were real translation sites, we added the translational inhibitor puromycin. As expected, this caused the frankenbody to disperse from all single mRNA sites within seconds (Fig. 5C, Movie S2). This confirms the HA frankenbody can indeed bind nascent peptide chains co-translationally.

**Figure 5.**
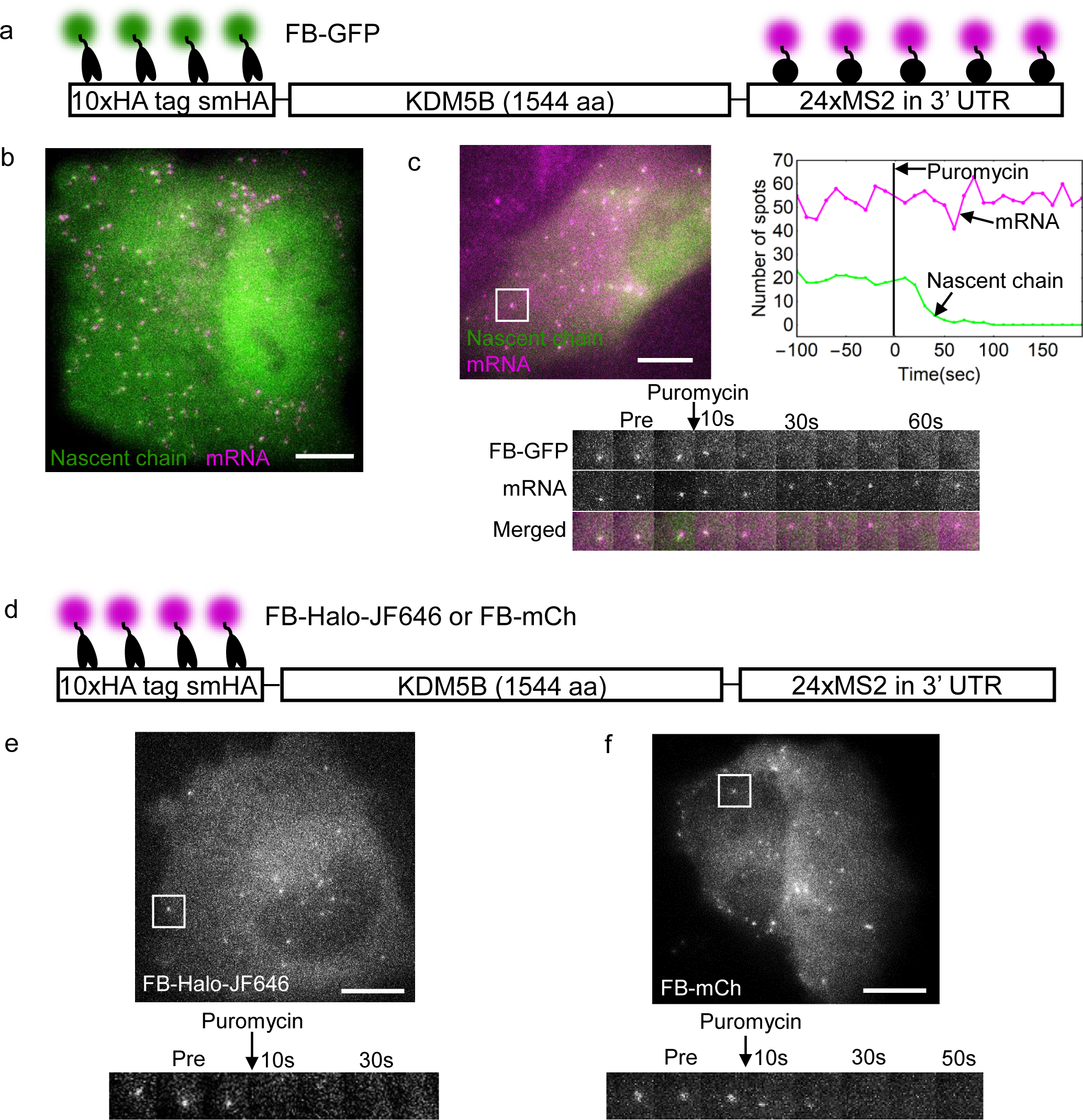
Tracking single mRNA translation in living U2OS cells with the HA frankenbody. (a) A diagram depicting frankenbody (FB-GFP; green) and MCP-HaloTag-JF646 (magenta) labeling HA epitopes and mRNA stem loops, respectively, in a KDM5B translation reporter. (b) A representative cell showing colocalization of FB-GFP (green) with KDM5B mRNA (magenta). (c) A representative cell (upper-left) showing the disappearance of nascent chain spots labeled by FB-GFP within seconds of adding the translational inhibitor puromycin. Upper-right: The number of nascent chain spots decreases while mRNA levels remain constant. Lower: a sample single mRNA montage. (d) A diagram depicting FB-Halo-JF646 or FB-mCh labeling HA epitopes in a KDM5B translation reporter. (e, f) Representative cells and single mRNA montages showing the loss of nascent chain spots labeled by (e) FB-Halo-JF646 or (f) FB-mCh upon puromycin treatment. Scale bars, 10 µm.

To ensure the frankenbody can also light up translation sites in multiple colors, we repeated experiments, but now using our other frankenbody constructs. For this, we co-transfected cells with either our mCherry or HaloTag frankenbody plasmids (FB-mCh or FB-Halo), together with our KDM5B translation reporter (Fig. 5D). In both cases, we could easily detect bright translation sites that responded to puromycin treatment (Fig. 5E and 5F, Movie S3 and S4). Collectively, these data demonstrate the HA frankenbody can be used to image translation in three colors spanning the imaging spectrum, demonstrating its potential and flexibility in multicolor experiments.

### Multiplexed imaging of single mRNA translation dynamics in living U2OS cells

With the ability to image translation in more than one color, we combined the HA frankenbody with the SunTag imaging system to simultaneously quantify the translation kinetics of two distinct mRNA species co-expressed in single living cells. Previously, the SunTag scFv has been fused to GFP (Sun-GFP) to monitor translation^18–21^. We therefore coupled this probe^23^ (after removing its HA epitope) with our complementary mCherry-tagged frankenbody probe (FB-mCh) (Fig. 6A). Co-transfecting these into living U2OS cells together with plasmids encoding SunTag-kif18b and smHA-KDM5B, we observed two distinct types of translation sites, those labeled entirely green by Sun-GFP and those labeled entirely magenta by frankenbody (Fig. 6B, Movie S5). After co-tracking hundreds of these translation sites, we quantified their mobilities. This revealed they both move with a similar kinetic, having a diffusion coefficient of 0.024±0.003 µm^2^/sec for Sun-GFP and 0.019±0.001 µm^2^/sec for FB-mCh. The similarity of their movement despite their different sequences shows that different mRNA types can nonetheless be translated in similar micro-environments.

**Figure 6.**
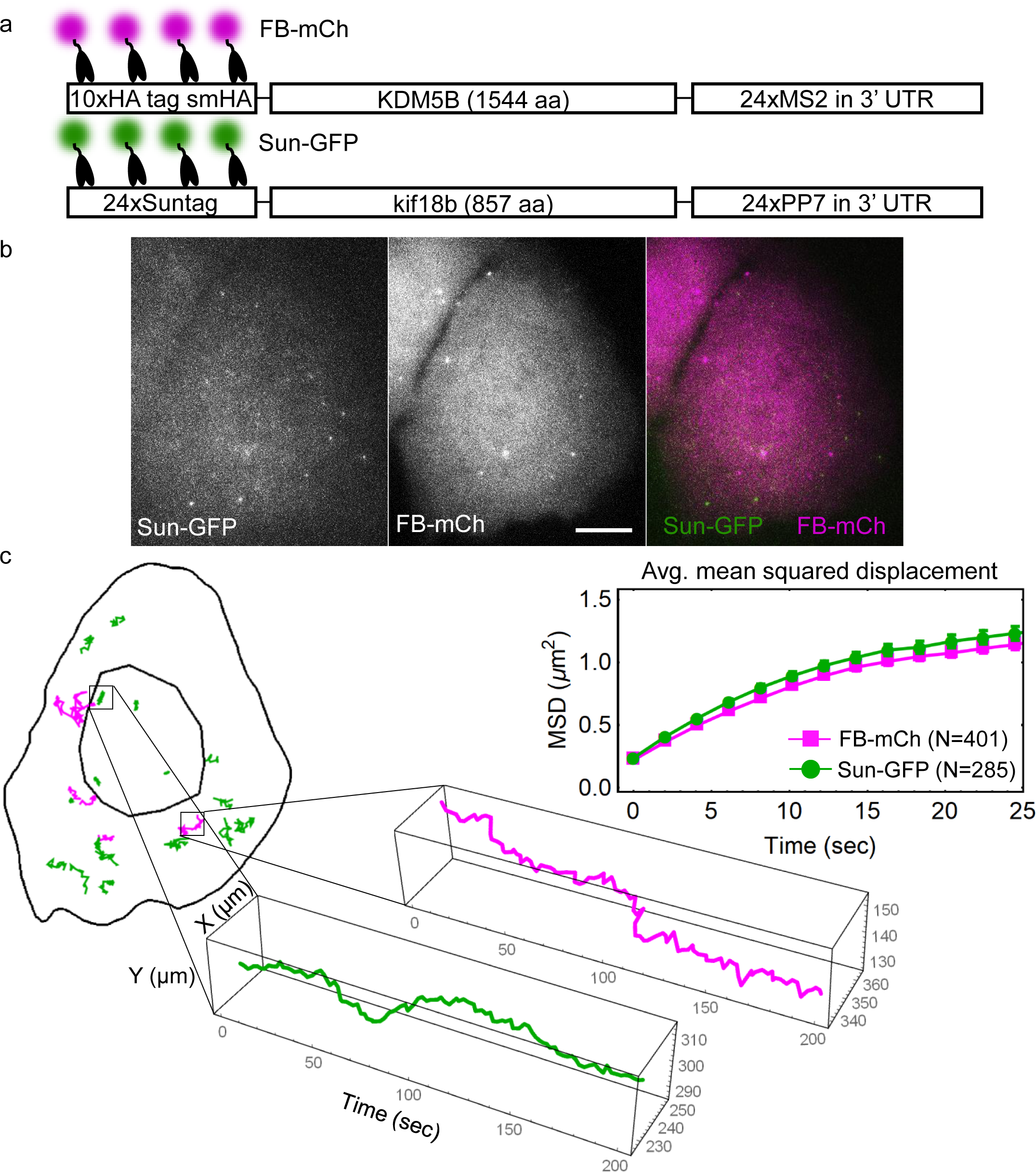
Multiplexed imaging of single mRNA translation dynamics in living U2OS cells. (a) A diagram depicting frankenbody (FB-mCh) and Sun-GFP labeling epitopes in the KDM5B and kif18b translation reporter constructs, respectively. (b) A representative living U2OS cell showing nascent chain translation spots labeled by FB-mCh (magenta) or Sun-GFP (green). (c) The average mean squared displacement of FB-mCh (magenta) and Sun-GFP (green) translation sites (upper-right). Error bars, SEM. Fits from the MSD curves show the diffusion coefficients are: 0.019±0.001 (90% CI) for FB-mCh and 0.024±0.003 (90% CI) for Sun-GFP. The mask and tracks of the cell in (b), and dynamics of a representative translation spot for each probe (Sun-GFP, green; FB-mCh, magenta). Scale bars, 10 µm.

Since both translation sites were labeled in different colors, we next wondered if they ever co-localized. The observation of colocalized translation sites would provide further evidence for multi-RNA “translation factories,” which our group^17^ and another^21^ have recently observed. In particular, we showed that approximately 5% of our smHA-KDM5B mRNA reporter is within multi-RNA factories^17^. Although we looked, we were unable to detect co-localized magenta and green translation sites. Since the smHA-KDM5B mRNA has different 3’ and 5’ untranslated regions (UTRs) as well as a different open reading frame (ORF) than SunTag-kif18b mRNA, while the translation reporters, used for the colocalization observation in our previous study, share identical 3’ and 5’ UTRs, but slightly different ORFs, these suggest that the composition of factories may be dictated in part through mRNA sequence elements.

### Monitoring local translation in living neurons with the HA frankenbody

Local translation is implicated in neuronal plasticity, memory formation, and disease^42^. The ability to image local translation at the single molecule level in living neurons would therefore be a valuable research tool to better understand these processes. To facilitate this, we tested if the HA frankenbody could be used to monitor single mRNA translation in living primary rat cortical neurons. When we co-transfected these cells with our KDM5B reporter^17^ and the GFP-tagged HA frankenbody (Fig. 7A), we were able to see distinct bright spots that diffused throughout the cell cytoplasm (Fig. 7B, Movie S6), reminiscent of the translation sites we had observed in U2OS cells. Again, we confirmed these were indeed translation sites by adding the translational inhibitor puromycin (Fig. 7C, Movie S7). Just seconds after the drug was added, the bright spots disappeared, just as they had in U2OS cells.

Unlike the mobility of mRNA we observed in U2OS cells, mRNA within neuronal dendrites displayed obvious directed motion events. For example, we regularly saw mRNA zip along linear paths within dendrites, achieving rapid translocations with rapid retrograde and anterograde transport over large distances up to 8 microns (Fig. 7D, Movie S6). The strong frankenbody signal within these fast-moving sites suggest translation is still active, despite the motored movement. These data therefore provide further support for a model in which translation is not necessarily repressed during trafficking^20^.

**Figure 7.**
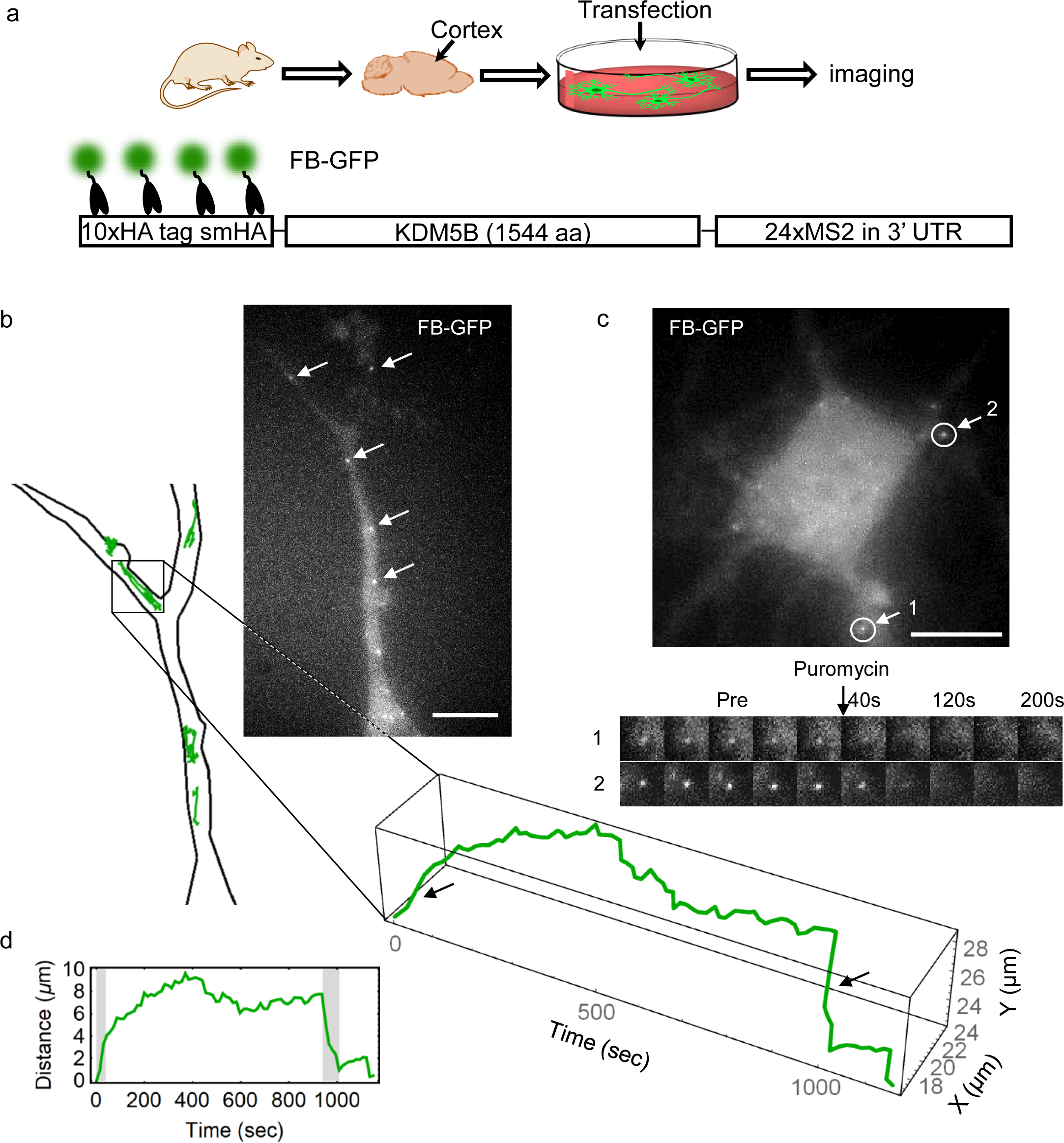
Monitoring local translation in living neurons with the HA frankenbody. (a) A diagram depicting the preparation of rat primary cortical neurons for imaging. (b) The dendrite of a sample living neuron expressing frankenbody (FB-GFP) and the smHA-KDM5B translation reporter. White arrows indicate translation sites that were tracked, as depicted in the cartoon below. The spatiotemporal evolution of one mRNA track with directed motion is shown through time. (c) A neuron just before puromycin was added. Two circled translation sites were tracked as they disappeared following the addition of puromycin. (d) The travel distance through time for the translation spot highlighted in b. Gray highlights in d and black arrows in b indicate directed motion events. Scale bars, 10 µm.

### Monitoring zebrafish development with the HA frankenbody

Arguably the most demanding types of imaging applications are in whole living animals. To verify that the HA frankenbody can also be applied in this way, we used it to monitor development in zebrafish embryos. The environment within embryos is complex and contains many potential non-specific binding targets. To express HA frankenbody in this complex environment, we microinjected mRNA encoding GFP-tagged frankenbody (FB-GFP) and HA-mCh-H2B into the yolk of one-cell stage zebrafish eggs. With this setup, HA frankenbody is expressed (i.e. translated) immediately without having to wait for the onset of transcription after the maternal-zygotic transition^43^. Following the initial injection, we co-loaded a positive control Fab (Cy5 conjugated) which specifically binds and lights up endogenous histone acetylation (H3K9ac) in the cell nucleus^25^ (Fig. S3A).

We began imaging embryo development with the HA frankenbody around the four- or eight-cell stage. At all timepoints we could see colocalization of the frankenbody with the HA-mCh-H2B target (Fig. 8A, Movie S8), although at earlier timepoints the concentrations of both were lower and therefore marked the nuclei only dimly compared to the positive control Fab. Nevertheless, we could detect all three signals in the nuclei of single mother and daughter cells throughout the entire 80-minute imaging time course (Fig. 8B, upper). Moreover, the signal from the frankenbody in the nucleus tightly correlated with the target HA-mCh-H2B signal, increasing steadily from a nuclear-to-cytoplasmic ratio of one to nearly 2.5 (Fig. 8B, lower). This single-cell trend was also observed in the population of cells (Fig. 8C). As a negative control, we repeated experiments in zebrafish embryos lacking target HA-mCh-H2B. In this case, the frankenbody was evenly distributed throughout the embryo (Fig. S3B, C, Movie S9), displaying a nuclear-to-cytoplasmic ratio close to one at all times. This confirms the HA frankenbody binds the HA epitope selectively and tightly *in vivo*, so it can be used to accurately monitor the concentration of target HA-tagged proteins in living organisms.

**Figure 8.**
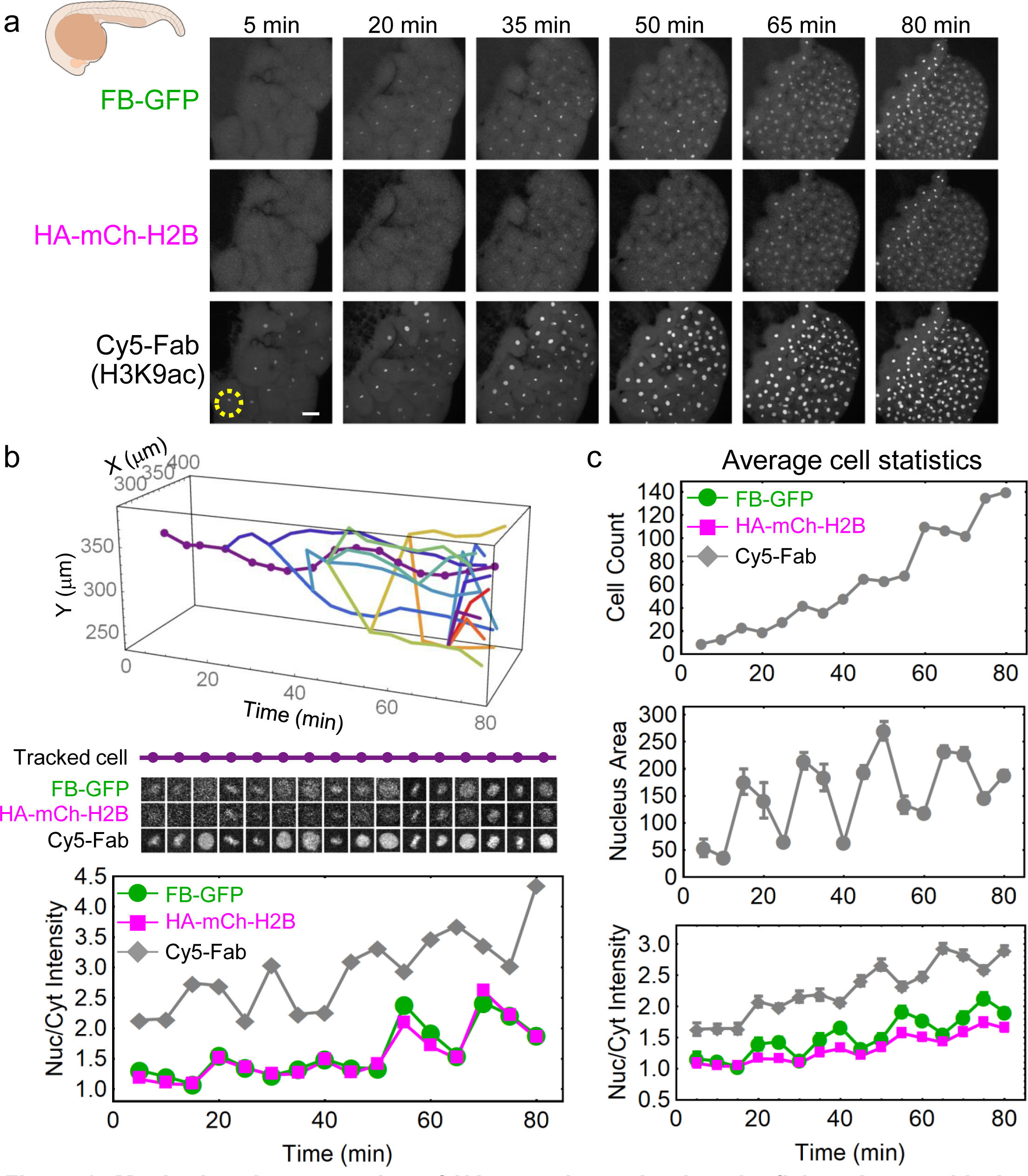
Monitoring the expression of HA-tagged proteins in zebrafish embryos with the HA frankenbody. (a) Max-projection images from a zebrafish embryo with frankenbody (FB-GFP) and HA-mCh-H2B. Cy5-Fab labels nuclear histone acetylation as a positive control. (b) A nuclei (dash circle in A) and its progeny tracked in development. The nuclear to cytoplasmic ratio (Nuc/Cyt) from FB-GFP (green circles), HA-H2B-mCh (magenta squares) and Fab (gray diamonds) in the parental nuclei (dotted line). (c) Cell count (top), average nuclear area (middle) and Nuc/Cyt ratios for all tracked nuclei. Error bars, SEM. Scale bar, 50 μm

## DISCUSSION

While scFvs have great potential for live-cell imaging, so far there are just a few documented examples of scFvs that fold and function in the reducing environment of living cells. Here we employed CDR loop grafting to generate with a 40% success rate stable and functional scFvs that bind the classic linear HA epitope tag *in vivo*. The resulting HA frankenbody is capable of labeling HA-tagged nuclear, cytoplasmic, and membrane proteins in multiple colors and in a diverse range of cellular environments, including primary rat cortical neurons and zebrafish embryos.

A major advantage of the HA frankenbody is it can be used to image single mRNA translation dynamics in living cells. This is because the target HA epitope (YPYDVPDYA, 9 aa) is small and linear. It therefore emerges quickly from the ribosome, so it can be co-translationally labeled by frankenbody almost immediately. It is also short enough to repeat many times in a single tag for signal amplification, as in the HA spaghetti monster tag^22^. In principle, epitopes could even be made conditionally accessible within a protein to monitor conformational changes. In contrast, almost all other antibody-based probes bind to 3D epitopes that span a large length of linear sequence space. In general, 3D epitopes take a relatively long time to be translated and fold before they become accessible for probe binding. Furthermore, they are too big to repeat more than a few times, so fluorescence is difficult to amplify to the degree necessary for single molecule tracking^22,23^.

Here we used the HA frankenbody to image single mRNA translation in both living U2OS cells and primary neurons. Unlike Fab, which cause neurons to peel during the loading procedure, HA frankenbody can be expressed in neurons without issue via transfection. We exploited this to demonstrate the mobility of translating mRNA is cell-type dependent. While our KDM5B translation reporter mRNA displayed largely non-directional, diffusive movement in U2OS cells, in neurons they were often motored. Neurons can be notoriously long, so motored mRNA movement provides a solution to the unique challenge of local protein production in distal neuronal dendrites and axons^44,45^. An open question is if translation is repressed during transport. On the one hand, certain mRNA are known to be actively repressed during trafficking^3,46^, perhaps to conserve energy. On the other hand, the Singer lab recently showed that in dendrites β-actin mRNA can be translated actively during transport^20^. Here we see a similar phenomenon, with our KDM5B translation reporter being rapidly motored over distances up to 8 microns in dendrites while retaining a strong translation signal. As our KDM5B reporter encodes the β-actin 3’ and 5’ UTR, our data demonstrate that the motored transport of translating β-actin transcripts within neurons is controlled by the 3’ or 5’ UTRs rather than the ORF.

Besides the HA frankenbody, there is only one other scFv capable of imaging single mRNA translation dynamics: the SunTag scFv^18,20,21,27,28^. Compared to the SunTag, the HA frankenbody binds it target epitope with lower affinity (nM versus pM). However, the SunTag epitope (EELLSKNYHLENEVARLKK, 19 aa) is over twice the length of the HA epitope^23^. Furthermore, the relatively new SunTag is not as ubiquitous as the HA-tag, which has enjoyed widespread use in the biomedical sciences for over thirty years^32^.

While the HA frankenbody and SunTag each have unique advantages, their combination creates a powerful genetically-encoded toolset to quantify single mRNA translation dynamics in two complementary colors. For example, their combination makes it possible to combine HA- and SunTag-epitopes in single mRNA reporters and thereby examine more than one open reading frame at a time or to create a gradient of translation colors for multiplexed imaging. In this study, we combined the HA frankenbody with the SunTag to quantify the spatiotemporal dynamics of two distinct mRNA species with different UTRs and ORFs. Unlike our earlier work with two mRNA species sharing common UTRs, in this case we did not observe colocalization of translation sites, i.e. “translation factories.” This would suggest that specific sequences within mRNAs likely dictate which translation factories they are recruited to, if any.

The genetic encodability of the HA frankenbody makes it a great value to researchers who have so far had to rely on expensive full-length antibodies purified from hybridoma cells to label HA-tagged proteins of interest. The HA frankenbody will therefore have an immediate and positive impact on the large cadre of researchers already employing the HA-tag in their studies, both *in vitro* and *in vivo*. With the HA frankenbody, researchers can simply transfect cells or animals expressing HA-tagged proteins with DNA or mRNA encoding the frankenbody fused to a fluorescent protein. This enables the visualization and quantification of HA-tagged protein expression, localization, and dynamics in living systems, not possible with traditional techniques and at a fraction of the cost. Here we demonstrated this potential by imaging the expression of HA-epitopes during the early stages of zebrafish embryo development. Even in this challenging environment, we were able to accurately detect and quantify HA-epitope expression in single cells and their progeny. In fact, the signal-to-noise from the frankenbody actually surpassed that of its target HA-H2B-mCh (Fig. 8B, C). This means the HA-frankenbody can be used to sensitively detect, amplify, and quantify the expression of short-lived proteins *in vivo*, similar to how the GFP-nanobody has been used to both amplify GFP fluorescence^13,14^ and image the spatiotemporal dynamics of short-lived, GFP-tagged transcription factors during Drosophila development^5^. Finally, the frankenbody can be genetically fused to other protein motifs to create a wide array of tools for live-cell imaging or manipulation of HA-tagged proteins, or it can be co-evolved with the HA-epitope into other complementary probe/epitope pairs via directed evolution techniques^47,48^. As shown in Fig. S1, the sequences of the two successful HA-scFv variants (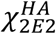 and 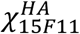) are slightly different, but their performance in living cells is almost identical according to our initial screen. Thus, many positions of the frankenbody can be mutated without destroying its functionality, demonstrating great potential for further evolution. We therefore anticipate the HA frankenbody will be a useful new imaging reagent to complement the growing arsenal of live-cell antibody-based probes^23,49–51^.

Given our success with CDR grafting in this study, in principle it should be relatively straightforward to generate more scFvs that bind additional targets besides the HA epitope tag. However, the functionality of CDR grafted scFvs is still difficult to predict^30^, so it remains unclear how generalizable the method is. According to our initial screen of scaffolds for frankenbodies, scFvs that have similar sequences will generally have a higher chance of being compatible grafting partners. We are therefore optimistic that as the price of determining antibody sequences continues to decrease, more scFvs will be constructed, tested and verified to fold and function in living cells. The availability of compatible grafting partners will therefore only increase, meaning less effort will be required to generate additional frankenbodies in the future. Ultimately, we envision a panel of complementary frankenbodies will be available to image in multiple colors the full lifecycles of proteins *in vivo* with high spatiotemporal resolution.

## Supporting information

## ACKNOWLEDGMENTS

We thank all members of the Stasevich lab for input and helpful suggestions, especially Kenneth Lyon for removing the HA epitopes from the Sun-GFP construct (Addgene #60907) and Lara Perinet for help with plasmid preparation. We also thank members of the CSU TagTeam; In particular, Dr. Chris Snow, Dr. Brian Geiss, and Steve Foster, for valuable discussions. Finally, we thank Laurie S. Minamide and Dr. James R. Bamburg of CSU for supplying neurons in this study.

## AUTHOR CONTRIBUTIONS

Conceptualization, N.Z. and T.J.S.; Methodology, N.Z., T.M., P.D.F., K.K., H.O., Y.S., H.K., and T.J.S.; Software, T.M. and T.J.S.; Validation, N.Z., P.D.F., K.K., and H.K.; Formal Analysis, N.Z. and T.J.S.; Investigation, N.Z., T.M., P.D.F., K.K., H.O., Y.S., H.K., and T.J.S.; Resources, N.Z., H.K., and T.J.S.; Data Curation, N.Z. and T.J.S.; Writing-Original Draft, N.Z. and T.J.S.; Writing-Review & Editing, N.Z. and T.J.S.; Visualization, N.Z., P.D.F., T.J.S., K.K., H.O., and H.K.; Supervision, T.J.S. and H.K.; Project Administration, N.Z. and T.J.S.; Funding Acquisition, T.J.S. and H.K.

## Data and materials availability

All data are included in the manuscript or supplemental materials and/or are available from the authors upon request.

## METHODS

### Plasmids Construction

Each chimeric anti-HA scFv tested in this study was constructed by grafting six CDR loops of an anti-HA antibody 12CA5 onto each selected scFv scaffold. The 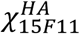 plasmid was constructed in two steps: 1) a CDR-loop grafted scFv gblock was synthesized *in vitro* by Integrated DNA Technologies (IDT) and ligated to a H4K20me1 mintbody 15F11 vector^35^ cut by EcoRI restriction sites via Gibson assembly (House prepared master mix); 2) the ^linker connecting scFv and EGFP, as well as EGFP, was replaced by a flexible (G^4^S)×5^ linker and the monomeric EGFP by Gibson assembly through NotI restriction sites. For the other 4 chimeric scFv plasmids, each CDR-loop grafted scFv gblock was synthesized *in vitro* and ligated into the 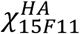 vector cut by EcoRI restriction sites via Gibson assembly.

The target plasmid HA-mCh-H2B was constructed in two steps: 1) the sfGFP of an Addgene plasmid sfGFP-H2B (Plasmid # 56367) was replaced with a 1×HA epitope tagged mCherry gblock synthesized *in vitro*, with a NotI restriction site inserted between the HA epitope and the mCherry coding sequence; 2) the 1×HA-epitope was replaced with a 4×HA-epitope gblock synthesized *in vitro* through AgeI and NotI restriction sites. Another target plasmid HA-mCh-β-actin was constructed by replacing the H2B with the β-actin encoding sequence via BglII and BamHI restriction sites. The β-actin was amplified from a published plasmid SM-β-actin^17^ using primers: 5’-TGG ACG AGC TGT ACA AGT CCG GAC TCA GAT CTG GAG GCT CCG GAG GCG ATG ATG ATA TCG CCG CGC TCG TCG TCG ACA ACG-3’; 5’-ATG ATC AGT TAT CTA GAT CCG GTG GAT CCC TTA GAA GCA TTT GCG GTG GAC GAT GGA GGG GCC GGA CT-3’. The smHA-Kv2.1 plasmid was generated by PCR amplification of the rat Kv2.1 coding sequence from pBK-Kv2.1^37^ and subsequent ligation into AsiSI and PmeI restriction sites on a published plasmid smHA-KDM5B^17^ using primers: 5’-CTG CAG GCG ATC GCC ACG AAG CAT GGC TCG CGC-3’; 5’-CGG GAG CAC TAG GGA TCA GAG TAT CGT TTA AAC GCT AGC T-3’. The plasmid smHA-H2B was constructed by ligating H2B cut from a published plasmid SM-H2B^17^ into a smHA-KDM5B^17^ intermediate plasmid through PstI and NheI restriction sites. mCh-H2B was generated by replacing sfGFP of the Addgene plasmid (Plasmid #56367) with a mCherry gblock synthesized *in vitro*.

The plasmids FB-mCh, FB-Halo, FB-SNAP were built by replacing the mEGFP of 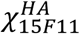 with mCherry, HaloTag and SNAP-tag gblocks synthesized *in vitro* respectively.

pET23b-FB-GFP, the plasmid for recombinant FB expression and purification, was generated by assembling a FB-GFP gene with a previously built plasmid pET23b-Sso7d^52,53^ by NdeI and NotI restriction sites. The FB-GFP encoding sequence was amplified from 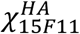 by PCR with primers: 5’-TTT GTT TAA CTT TAA GAA GGA GAT ATA CAT ATG GCC GAG GTG AAG CTG GTG GAG-3’; 5’-TGG TGG TGC TCG AGT GCG GCC GCC TTG TAC AGC TCG TCC ATG CCG AGA G-3’.

For the translation assay, the reporter construct smHA-KDM5B-MS2 was constructed previously^17^. The SunTag-kif18b reporter plasmid was purchased from Addgene (Plasmid # 74928), and its scFv plasmid (Plasmid #60907) was modified by removing the HA epitope encoded in the linker. The HA epitope was removed by site-directed mutagenesis with QuikChange Lightning (Agilent Technologies) per the manufacturer’s instruction using primers: 5′-CCT CCG CCT CCA CCA GCG TAA TCT GAA CTA GCG GTT CTG CCG CTG CTC ACG GTC ACC AGG GTG CCC −3′; 5′-GGG CAC CCT GGT GAC CGT GAG CAG CGG CAG AAC CGC TAG TTC AGA TTA CGC TGG TGG AGG CGG AGG −3′.

The gblocks were synthesized by Integrated DNA Technologies and the recombinant plasmids were sequence verified by Quintara Biosciences. All plasmids used for imaging translation were prepared by NucleoBond Xtra Midi EF kit (Macherey-Nagel) with a final concentration about 1 mg/mL.

### U2OS Cell Culture, Transfection and Bead Loading

U2OS cells were grown in an incubator at 37°C, humidified, with 5% CO_2_ in DMEM medium (Thermo Scientific) supplemented with 10% (v/v) fetal bovine serum (Altas Biologicals), 1mM L-glutamine (Gibco) and 1% (v/v) penicillin-streptomycin (Gibco or Invitrogen).

For tracking protein localization, cells were plated into a 35 mm MatTek chamber (MatTek) 2 days before imaging and were transiently transfected with Lipofectamine^TM^ LTX reagent with PLUS reagent (Invitrogen) according to the manufacturer’s instruction 18~24 hours prior to imaging.

For imaging translation with mRNA labeled by MCP-Halo protein, cells were plated into a 35 mm MatTek chamber (MatTek) the day before imaging. On the day of imaging, before bead loading, the medium in the MatTek chamber was changed to Opti-MEM (Thermo Scientific) with 10% fetal bovine serum. A mixture of plasmids (smHA-KDM5B/FB) and purified MCP-HaloTag protein^17^ were bead loaded as previously described^6,17,26^. Briefly, after removing the Opti-medium from the MatTek chamber, 4 uL of a mixture of plasmids (1 ug each) and MCP-HaloTag (130 ng) in PBS was pipetted on top of the cells and ~106 µm glass beads (Sigma Aldrich) were evenly distribute on top. The chamber was then tapped firmly 7 times, and Opti-medium was added back to the cells. 3 hours after bead loading, the cells were stained in 1 mL of 0.2 nM of JF646-HaloTag ligand^41^ diluted in phenol-red-free complete DMEM. After a 20 min incubation, the cells were washed three times in phenol-red-free complete DMEM to remove glass beads and unliganded dyes. The cells were then ready for imaging.

For imaging translation without mRNA labeling, the cells plated on MatTek chamber were transiently transfected with the plasmids needed, smHA-KDM5B/FB, and/or SunTag-kif18b/Sun (with the HA epitope removed), with Lipofectamine^TM^ LTX reagent with the PLUS reagent (Invitrogen) according to the manufacturer’s instruction on the day of imaging. 3 hours post transfection, the medium was changed to phenol-red-free complete DMEM. The cells were then ready for imaging.

### Purification of FB-GFP

*E. coli* BL21 (DE3) pLysS cells transformed with pET23b-FB-GFP were grown at 37°C to a density of OD600 0.6 in 2xYT medium containing Ampicillin (100 mg/L) and Chloramphenicol (25 mg/L) with shaking. Isopropyl-β-D-thiogalactoside (IPTG) was added to induce protein expression at a final concentration of 0.4 mM, and the temperature was lowered to 18°C. Cells were harvested after 16 hours by centrifugation and resuspended in PBS buffer supplemented with 300 mM NaCl, protease inhibitors (ThermoFisher), 0.2 mM AEBSF (20 ml/1L culture) and lysed by sonication. Lysate was clarified by centrifugation. The supernatant was loaded onto 2 connected HisTrap HP 5 ml columns (GE Healthcare), washed and eluted by a linear gradient of 0-500 mM imidazole. The fractions containing the protein of interest were pooled, concentrated using Amicon Ultra-15 30 kDa MWCO centrifugal filter unit (EMD Millipore) and loaded onto a size-exclusion HiLoad Superdex 200 PG column (GE healthcare) in HEPES-based buffer (25 mM HEPES pH 7.9, 12.5 mM MgCl_2_, 100 mM KCl, 0.1 mM EDTA, 0.01 % NP40, 10% glycerol and 1 mM DTT). The fractions containing FB-GFP protein were collected, concentrated, and stored at −80°C after flash freezing by liquid nitrogen.

### Immunostaining

U2OS cells were transiently transfected with HA-mCh-H2B or HA-mCh-β-actin with Lipofectamine^TM^ LTX reagent with the PLUS reagent (Invitrogen). 26 hours post transfection, the cells were fixed with 4% paraformaldehyde (Electron Microscopy Sciences) for 10 minutes at room temperature, permeabilized in 1% Triton 100 in PBS, pH 7.4 for 20 minutes, blocked in Blocking One-P (Nacalai Tesque) for 20 minutes, and stained at 4°C overnight with purified FB-GFP protein (0.5 ug/mL in 10% blocking buffer). The next morning, the cells were washed with PBS, and the protein of interest was imaged by an Olympus IX81 spinning disk confocal (CSU22 head) microscope using a 100x oil immersion objective (NA 1.40) under the following conditions: 488 nm and 561 nm sequential imaging for 50 timepoints without delay, 20% laser power, 2×2 spin rate, 100 ms exposure time. Images were acquired with a Photometrics Cascade II CCD camera using SlideBook software (Intelligent Imaging Innovations). The immunostaining images were generated by averaging 50 timepoint images for each channel by Fiji^54^.

### Western Blots

U2OS cells were transiently transfected with HA-mCh-H2B or HA-mCh-β-actin with Lipofectamine^TM^ LTX reagent with the PLUS reagent. 24 hours post transfection, the cells were lysed in RIPA buffer with cOmplete Protease Inhibitor (Roche). 10 uL of each cell lysate was loaded on a NuPAGE^TM^ 4%~12% Bis-Tris protein gel (Invitrogen) and run for 60 minutes at 100 V and 25 minutes at 200 V. Proteins were transferred to a PVDF membrane (Invitrogen), blocked in blocking buffer (5% milk powder in 0.05% TBS-Tween 20) for 1 hour, and stained for 1 hour with either purified FB-GFP protein (0.5ug/mL in blocking buffer) or overnight with anti-HA parental antibody 12CA5 (Sigma-Aldrich, 0.5ug/mL in blocking buffer). For the primary antibody 12CA5, an additional incubation for 1 hour with anti-mouse antibody/HRP (5000-fold dilution in blocking buffer) was done. For FB-GFP, the protein of interest was detected from the GFP fluorescence using a Typhoon FLA 9500 (GE Healthcare Life Sciences) with the following conditions: excitation wavelength 473 nm, LPB filter (≥510 nm), 300 V photomultiplier tube and 10 µm pixel size. For HRP conjugated antibody, the protein of interest was detected by chemiluminescent ECL Western Blotting substrate (Pierce) according to the manufacturer’s instruction.

### Fluorescence Recovery after Photobleaching (FRAP)

To study the binding affinity of FB to HA epitopes in living cells, FRAP experiments were performed on cells transiently transfected with HA-mCh-H2B (1.25ug) and FB-GFP (1.25ug) 24 hours before FRAP. The images were acquired using an Olympus IX81 spinning disk confocal (CSU22 head) microscope coupled to a Phasor photomanipulation unit (Intelligent Imaging Innovations) with a 100x oil immersion objective (NA 1.40). Before photobleaching, 20 frames were acquired with 1 s time interval. The images were captured using 488 nm laser with 100 ms exposure time at 20% laser power followed by 561 nm laser with 15 ms exposure time at 20% laser power. The spinning disk was set up at 1×1 spin rate. After acquiring 20 pre-FRAP images, the 488 laser with 100% laser power and a 100 ms exposure time was used to photobleach a circular region in the nucleus. After photobleaching, 30 images were captured without a time interval delay, and then 100 images with a 1 s time interval delay and 100 images with a 5 s time interval delay were acquired using the same imaging settings as the pre-FRAP images. The fluorescent intensity through time of the photobleached spot were exported using the Slidebook software. The fluorescent intensity of the nucleus and background were obtained by Fiji^54^ after correcting for cell movement using the StackReg Fiji plugin^55^. The FRAP curve and t_half_ were obtained using easyFRAP-web56, according to the website instructions.

### MCP purification

MCP-HaloTag was purified as described previously^17^. Briefly, the His-tagged MCP-HaloTag was purified through a Ni-NTA-agarose (Qiagen) packed column per the manufacturer’s instructions, with minor modifications. *E. coli* expressing the interested protein was lysed in a PBS buffer with a complete set of protease inhibitors (Roche) and 10 mM imidazole. The resin was washed with PBS-based buffer containing 20 and 50 mM imidazole. The protein was then eluted in a PBS buffer with 300 mM imidazole. The eluted His-tagged MCP was dialyzed in a HEPES-based buffer (10% glycerol, 25 mM HEPES pH 7.9, 12.5 mM MgCl2, 100 mM KCl, 0.1 mM EDTA, 0.01 % NP-40 detergent, and 1 mM DTT), snap-frozen in liquid nitrogen, and then stored at −80°C.

### Nascent Chain Tracking

Nascent chain tracking was performed as described previously^17^. Briefly, the reporter plasmid smHA-KDM5B and the FB construct were either transiently transfected without the MCP-HaloTag protein or bead loaded with MCP-HaloTag protein into U2OS cells plated on a 35 mm MatTek chambers 4~6 hours before imaging. 3 hours later, if MCP-HaloTag protein was bead loaded, the cells were stained with the JF646-HaloTag ligand, then washed with phenol-red-free complete DMEM medium. If no MCP-HaloTag protein was needed, the medium of the cells was changed to phenol-red-free complete DMEM medium 3 hours post transfection. The cells were then ready for imaging.

For multiplexed imaging, 2 reporter plasmids, smHA-KDM5B and SunTag-kif18b, as well as 2 probes, FB-mCh and Sun-GFP (with the HA epitope removed), were transiently transfected into U2OS cells plated on a MatTek chamber 4~6 hours before imaging. 3 hours later, the medium of the cells was changed to phenol-red-free complete DMEM medium. The cells were then ready for imaging.

### Imaging condition for translation and colocalization assays

To image single mRNAs and their translation status with FB, a custom-built widefield fluorescence microscope based on an inclined illumination (HILO) scheme was used^17,57^. Briefly, the excitation beams, 488, 561, 637 nm solid-state lasers (Vortran), were coupled and focused off-axis on the rear focal plane of the objective lens (APON 60XOTIRF, Olympus). The emission signals were split by an imaging grade, ultra-flat dichroic mirror (T660lpxr, Chroma). The longer emission signals (far-red) after splitting were passed through a bandpass filter (FF01-731/137-25, Semrock). The shorter emission signals (red and green) after splitting were passed through either a bandpass filter for red (FF01-593/46-25, Semrock) or a bandpass filter for green (FF01-510/42-25, Semrock) installed in a filter wheel (HS-625 HSFW TTL, Finger Lakes Instrumentation). The longer (far-red) and the shorter (red and green) emission signals were detected by separate two EM-CCD cameras (iXon Ultra 888, Andor) by focusing with a 300 mm achromatic doublet lenses (AC254-300-A-ML, Thorlabs). The combination of 60× objective lens from Olympus, 300 mm tube lens, and iXon Ultra 888 produces 100× images with 130 nm/pixel. A stage top incubator for temperature (37 °C), humidity, and 5% CO_2_ (Okolab) is equipped on a piezoelectric stage (PZU-2150, Applied Scientific Instrumentation) for live cell imaging. The lasers, the cameras, the piezoelectric stage, and the filter wheel were synchronized by an open source micro controller, Arduino Mega board (Arduino). Imaging acquisition was performed using open source Micro-Manager software (1.4.22)^58^.

The imaging size was set to the center 512 × 512 pixels^2^ (66.6 x 66.6 µm^2^), and the camera integration time was set to 53.64 ms. The readout time of the cameras from the combination of our imaging size, readout mode (30 MHz), and vertical shift speed (1.13 µs) was 23.36 ms, resulting in our imaging rate of 13 Hz (70 ms per image). Red and green signals were imaged alternatively. The emission filter position was changed during the camera readout time. To minimize the bleed-through, the far-red signal was simultaneously imaged with the green signal. To capture the whole thickness of cells, 13 z-stacks with a step size of 500 nm (6 µm in total) were imaged using the piezoelectric stage. This resulted in our total cellular imaging rate of 1 Hz for imaging either red or green signals, and 0.5 Hz for imaging both red and green signals regardless of far-red imaging.

For Fig. 1D, 1E, 2A, and 2D, a single plane of the cells was imaged continuously at 6.5 Hz for 100 time points and averaged throughout the time. For Fig. 2 B, the cell was imaged continuously at 0.5 Hz with 13 z-stacks per timepoint and averaged throughout the time. The acquired averaged 13 z-stacks were deconvolved using Fiji. For Fig. 2C, the cells were imaged at a single time point with 13 z-stacks. For Fig. 5B, 5C, 5E, 5F and 7C, cells were imaged every 10 sec with 13 z-stacks per timepoint. For Fig. 6B, the cell was imaged continuously at 0.5 Hz with 13 z-stacks per timepoint. For Fig. 7B, the cells were imaged every 14 sec with 13 z-stacks per timepoint.

### Particle tracking

Single particle detection and tracking was performed on maximum intensity projection images with custom Mathematica (Wolfram Research) code, as previously described^17^. Briefly, the images were processed with a bandpass filter to highlight particles, and then binarized to detect their intensity-centroids as positions using the built-in Mathematica routine ComponentMeasurements. Detected particles were tracked and linked through time via a nearest neighbor search. The precise coordinates (super-resolved locations) of mRNAs and translation sites were determined by fitting (using the built-in Mathematica routine NonlinearModelFit) the original images to 2D Gaussians of the following form:

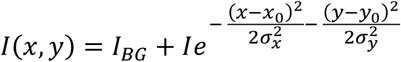

where *I*_*BG*_ is the background fluorescence, *I* the particle intensity, (*x*_0_, *y*_0_) the particle location, and (*σ*_*x*_, *σ*_*y*_) spreads of the particle. The offset between the two cameras was registered using the built-in Mathematica routine FindGeometricTransform to find the transform function that best aligned the fitted positions of 100 nm diameter Tetraspeck beads evenly spread out across the image field-of-view.

For Fig. 6C, average mean squared displacements were calculated from the Gaussian-fitted coordinates (from 2D maximum intensity projection images). The diffusion constant was obtained by fitting the first 10 time points to a line with slope *m* = 4*D*, where *D* is the diffusion coefficient.

### Puromycin Treatment

U2OS cells transiently transfected with smHA-KDM5B and FB or bead loaded with smHA-KDM5B, FB and MCP-HaloTag were imaged as above with 10 s intervals between frames. After acquiring 5 or 10 timepoints as pre-treatment images, cells were treated with a final concentration of 0.2 mg/mL puromycin right before acquiring the 6^th^ or 11^th^ timepoint. After puromycin was added, the cells were imaged under the same conditions used for the pre-treatment imaging until the translation spots disappeared.

### Neuron Culture and Transfection

Rat cortical neurons were obtained from the discarded cortices of embryonic day (E)18 fetuses which were previously dissected to obtain the hippocampus, and frozen in Neurobasal medium (ThermoFisher Scientific) containing 10% fetal bovine serum (FBS, Atlas Biologicals) and 10% DimethylSulfoxide (Sigma-Aldrich, D8418) in liquid nitrogen. Cryopreserved rat cortical neurons were plated at a density of ∼15,000–30,000 cells/cm^2^ on MatTek dishes (MatTek) and cultured in Neurobasal medium containing 2% B27 supplement (ThermoFisher Scientific), 2mM L-Alanine/L-Glutamine and 1% FBS (Atlas Biologicals). Transfections were performed after 5–7 days in culture by using Lipofectamine 2000 (ThermoFisher Scientific) according to the manufacturer’s instructions. Neurons co-expressing smHA-Kv2.1 and FB-GFP, or GFP-Kv2.1 were imaged 1-7 days post-transfection. For translation assays, neurons were imaged 4-12h post-transfection. All neuron imaging experiments were carried out in a temperature-controlled (37°C), humidified, 5% CO_2_ environment in Neurobasal medium without phenol red (ThermoFisher Scientific). Neuronal identity was confirmed by following processes emanating from the cell body to be imaged for hundreds of microns to ensure they were true neurites.

### Monitoring Zebrafish development

To visualize FB-GFP in zebrafish embryo, mRNAs for FB-GFP and HA-mCh-H2B were prepared. DNA fragments coding FB-GFP and HA-mCh-H2B were inserted into a plasmid containing the T7 promoter and poly A^59^. The subsequent plasmids (T7-FB-GFP and T7-HA-mCh-H2B) were linearized with the XbaI restriction site for *in vitro* transcription using mMESSAGE mMACHINE kit (ThermoFisher Scientific). RNA was purified using RNeasy Mini Elute Cleanup Kit (QIAGEN) and resuspended in water. Before microinjection, zebrafish (AB) eggs were dechorionated by soaking in 2 mg/ml pronase (Sigma Aldrich; P5147) in 0.03% sea salt for 10 minutes. A mixture (~0.5 nl) containing mRNA (200 pg each for FB-GFP and HA-mCh-H2B) was injected into the yolk (near the cell part) of 1-cell stage embryos. For a negative control, PBS was injected instead of HA-mCh-H2B mRNA. 5-10 minutes after mRNA injection, Cy5-labeled Fab specific to endogenous histone H3 Lys9 acetylation (CMA310)^25^ was injected (100 pg in ~0.5 nl). Injected embryos were incubated at 28°C until the 4-cell stage and embedded in 0.5% agarose (Sigma Aldrich, A0701) in 0.03% sea salt with the animal pole down on a 35-mm glass bottom dish (MatTek). The fluorescence images were collected using a confocal microscope (Olympus; FV1000) equipped with a heated stage (Tokai Hit) set at 28°C and a UPLSAPO 30x silicone oil immersion lens (NA 1.05), operated by the built-in FV1000 software FLUOVIEW ver.4.2. Three color images were sequentially acquired every 5 minutes using 488-, 543-, and 633-nm lasers (640 × 640 pixels; pinhole 800 μm; 2.0 μs/pixel) without averaging. Maximum intensity projections were created from 20 z-stacks with 5 μm.

Nuclei within zebrafish embryos were tracked in 4D using the Fiji plugin TrackMate^60^. Results were post-processed and plotted with *Mathematica*. To quantify the number and area of nuclei and the average nuclear, cytoplasmic, and nuclear:cytoplasmic intensity through time, the intensity of all nuclei in maximum intensity projections was measured using the built-in *Mathematica* function *ComponentMeasurements*. *ComponentMeasurements* requires binary masks of the objects to be measured. Binary masks of the nuclei were made using the built-in Mathematica function *Binarize* with an appropriate intensity threshold to highlight just nuclei in images from Cy5-labeled Fab (specific to endogenous histone H3 Lys9 acetylation). Masks of the cytoplasm around each nuclei were made by dilating the nuclear masks by 4 pixels (using the built-in command *Dilation*) and then subtracting from the dilated mask the original nuclear masks dilated by 1 pixel. This creates ring-like masks around each nuclei, from which the average cytoplasmic intensity was measured.

### Surface plasma resonance

Binding kinetics of purified FB to the HA epitope tag was measured by surface plasma resonance (OpenSPR, Nicoyalife). After biotin-labeled HA peptide was captured by a Streptavidin sensor chip (Nicoyalife), diluted purified FB-GFP in PBS running buffer, pH 7.4, was slowly flowed over the sensor chip for 5 min to allow interaction. The running buffer was then allowed to flow for 10 min to collect the dissociation data. The non-specific binding curve was obtained by flowing the same concentration of FB-GFP in the same running buffer over a different Streptavidin sensor chip (Nicoyalife). The data from the control was collected exactly as in the experiment. After subtracting the signal response against the no-HA peptide captured Streptavidin sensor chip, the signal response vs time curve was obtained, as shown in Fig. S2. Binding kinetic parameters were obtained by fitting the curve to a one-to-one binding model using TraceDrawer (Nicoyalife) software (Fig. S2).

---

**Figure S1.**
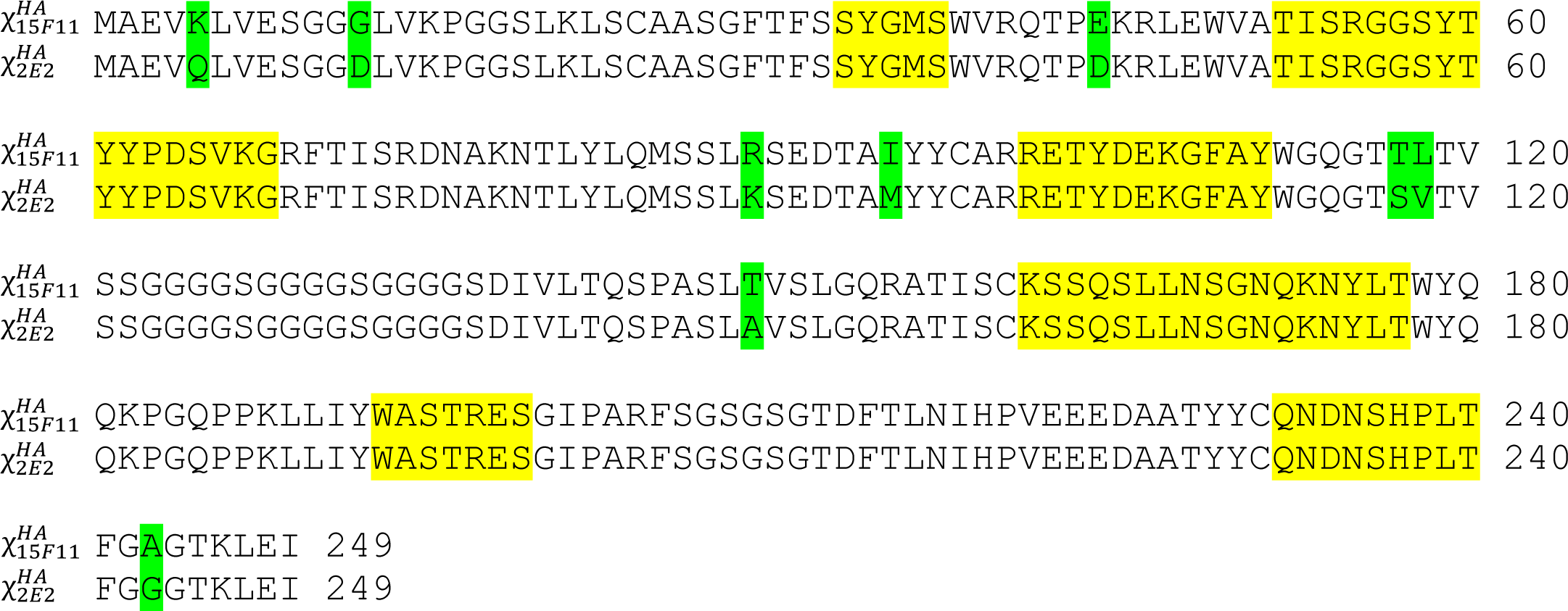
The sequence of frankenbodies. Six CDRs are highlighted in yellow. Mismatched amino acids are highlighted in green.

**Figure S2.**
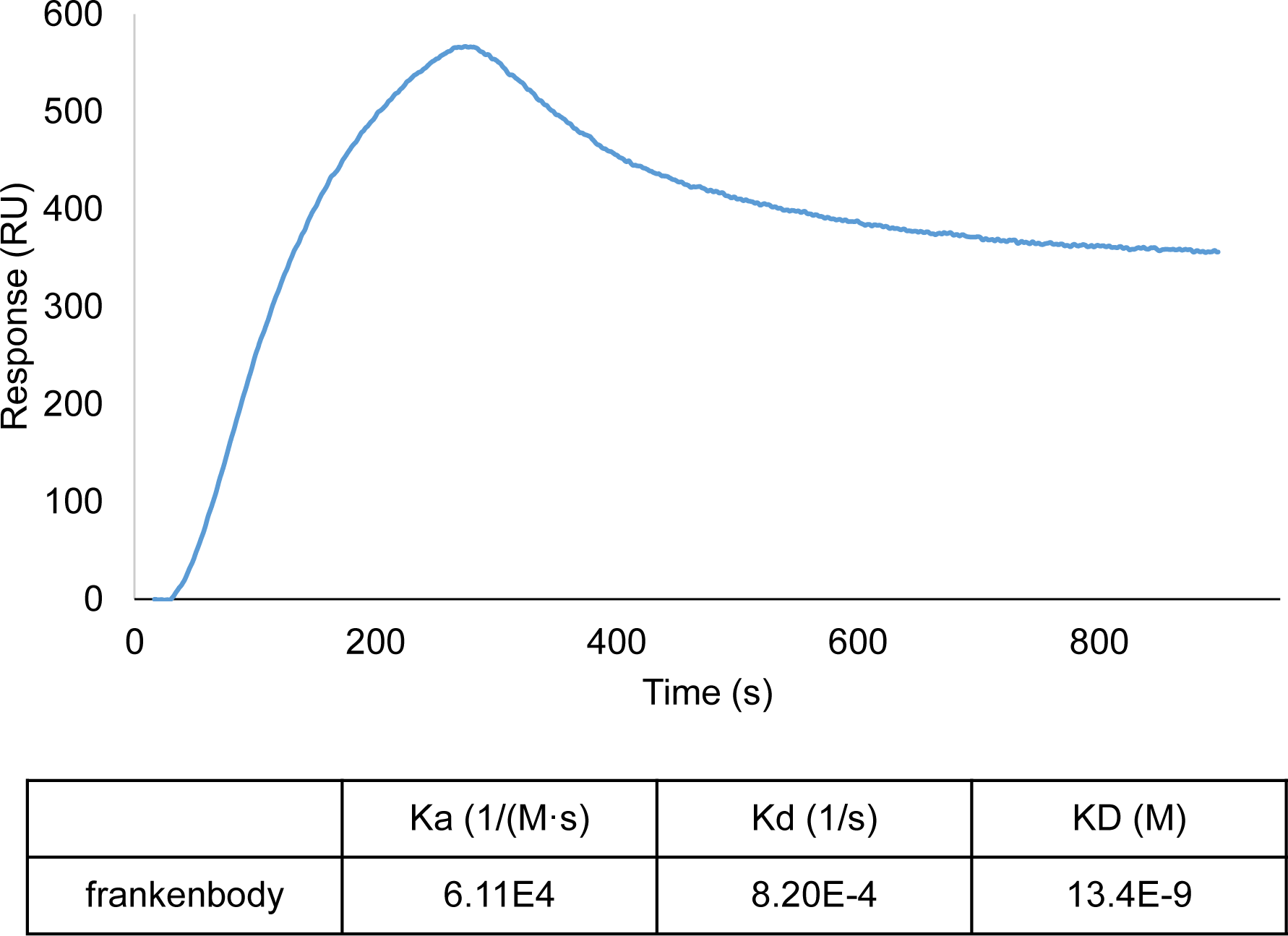
Binding kinetics of frankenbody to HA tag *in vitro*.

**Figure S3.**
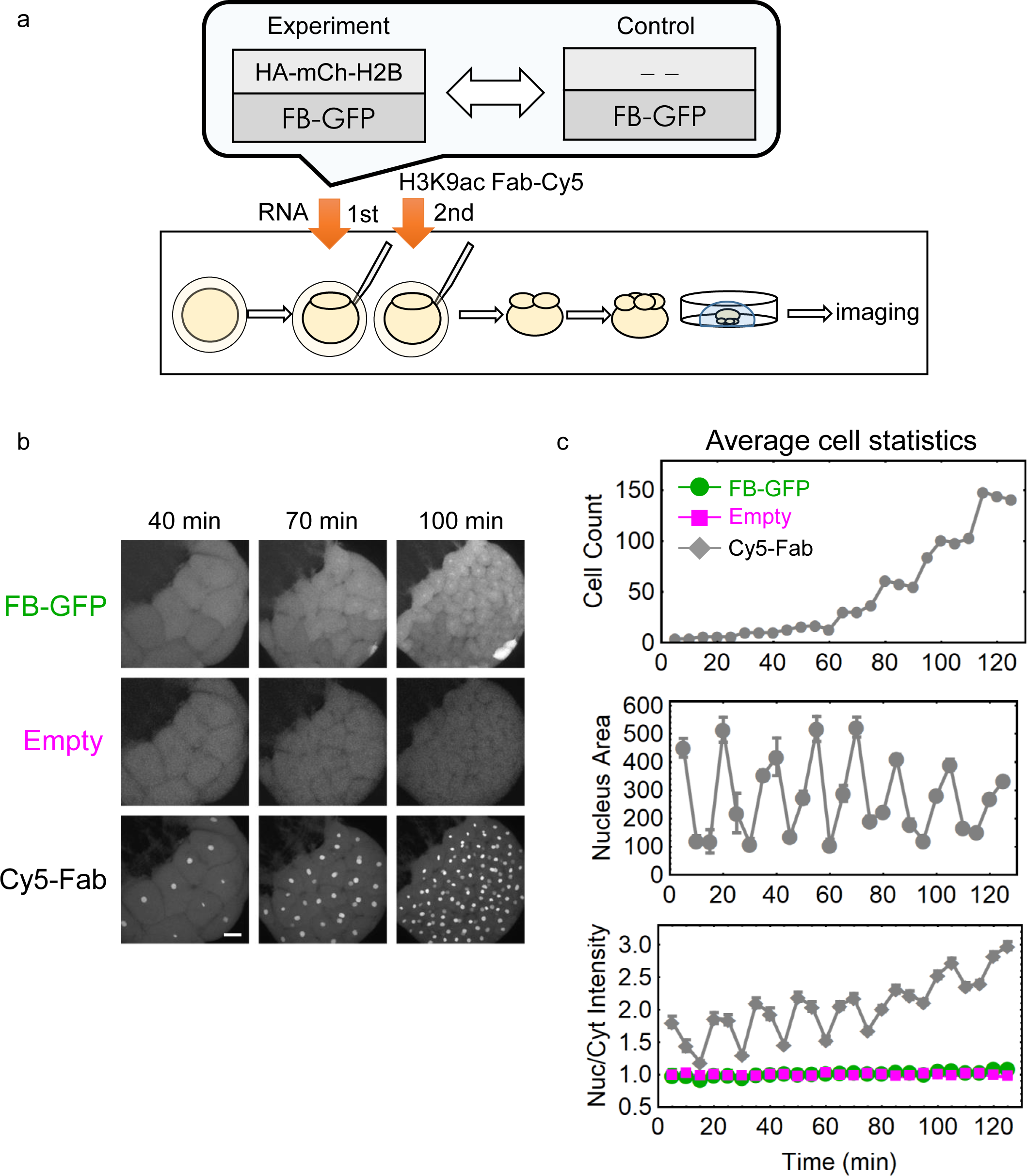
Related to Figure 8. HA frankenbody does not bind non-specifically in zebrafish embryos lacking HA epitopes. (a) A diagram depicting the timing of zebrafish embryo imaging experiments. (b) Sample max-projection images from a control zebrafish embryo with frankenbody (FB-GFP; green), but lacking target HA-mCh-H2B (Empty; magenta). Positive control Cy5-Fab marks histone acetylation in nuclei. (c) Cell count (top), average nuclear area (middle) and nuclear to cytoplasmic (Nuc/Cyt) ratio in all tracked nuclei through time. Error bars, SEM. Scale bar: 50 µm

## Supplemental movie legends

**Movie S1. Related to Figure 5B. Frankenbody tracks single mRNA translation in living U2OS cells.** Nascent chain was labeled by FB-GFP shown in green. mRNA was labeled by MCP-Halo-JF646 shown in magenta. Images were acquired every 10 sec (movie duration is 9 min 50 sec). Movie field of view is 66.56 × 66.56 µm.

**Movie S2. Related to Figure 5C. Puromycin treatment of FB-GFP tracking single mRNA translation in living U2OS cells.** Puromycin was added at 1 min 40 sec, right before frame 11. Images were acquired every 10 sec (movie duration is 4 min 50 sec). Movie field of view is 46.15 × 43.03 µm.

**Movie S3. Related to Figure 5E. Puromycin treatment of FB-Halo tracking single mRNA translation in living U2OS cells.** Puromycin was added at 1 min 40 sec, right before frame 11. Images were acquired every 10 sec (movie duration is 9 min 50 sec). Movie field of view is 66.56 × 66.56 µm.

**Movie S4. Related to Figure 5F. Puromycin treatment of FB-mCh tracking single mRNA translation in living U2OS cells.** Puromycin was added at 1 min 40 sec, right before frame 11. Images were acquired every 10 sec (movie duration is 9 min 50 sec). Movie field of view is 66.56 × 66.56 µm.

**Movie S5. Related to Figure 6B. Multiplexed imaging of single mRNA translation in living U2OS cells.** 2 orthogonal pairs of probe/translation reporter: Sun-GFP/SunTag-Kif18b (green) and FB-mCh/smHA-KDM5B (magenta) track single mRNA translation simultaneously in one living U2OS cell. Images were acquired every 2 sec (movie duration is 3 min 18 sec). Movie field of view is 66.56 × 66.56 µm.

**Movie S6. Related to Figure 7B. Frankenbody tracks single mRNA translation in living neurons.** A dendrite of a sample living neuron expressing frankenbody (FB-GFP) and the smHA-KDM5B translation reporter. Images were acquired every 14 sec (movie duration is 23 min 25 sec). Movie field of view is 66.56 × 66.56 µm.

**Movie S7. Related to Figure 7C. Puromycin treatment of FB-GFP tracking single mRNA translation in living neurons.** Puromycin was added at 3 min 20 sec, right before frame 6. Image was acquired every 40 sec (movie duration is 14 min). Movie field of view is 33.8 × 33.67 µm.

**Movie S8. Related to Figure 8A. HA frankenbody binds target epitopes in zebrafish embryos.** Single-channel movies showing the HA frankenbody binding specifically to target HA-epitopes in zebrafish embryos (Left top: FB-GFP; Right top: HA-mCh-H2B; Left bottom: Cy5-Fab). A merge of all 3 channels is also shown (Right bottom; FB-GFP: green; HA-mCh-H2B: red; Cy5-Fab: blue). Images were acquired every 5 min (movie duration is 80 min). Movie field of view is 423.68 × 423.68 µm.

**Movie S9. Related to Figure S3. HA frankenbody does not bind non-specifically in zebrafish embryos lacking HA epitopes.** Single-channel movies showing the HA frankenbody in zebrafish embryos lacking HA-epitopes (Left top: FB-GFP; Right top: empty; Left bottom: Cy5-Fab). A merge of all 3 channels is also shown (Right bottom; FB-GFP: green; empty: red; Cy5-Fab: blue). Images were acquired every 5 min (movie duration is 80 min). Movie field of view is 423.68 × 423.68 µm.

